# Distinct Auditory Thalamocortical Pathologies Underlie Emerging Neurophysiological Dysfunction in a Cln3 Mouse Model of Batten Disease

**DOI:** 10.64898/2026.06.01.729359

**Authors:** Yanya Ding, Jingyu Feng, Viollandi Prifti, Grace A. Rico, Alexander G. Solorzano, Hayley E. Chang, Samantha A. Spallina, Edward G. Freedman, John J. Foxe, Kuan Hong Wang

**Affiliations:** Department of Neuroscience, University of Rochester Medical Center, Rochester, NY, 14642, USA

**Keywords:** CLN3 disease, Lysosomal storage disorder, Subunit C of Mitochondrial ATP Synthase (SCMAS) Age- and sex-dependent differences, Auditory thalamocortical circuits, Auditory duration mismatch negativity (MMN) Auditory evoked potential (AEP)

## Abstract

CLN3 disease, the most common form of the Neuronal Ceroid Lipofuscinoses (NCLs), causes progressive cognitive decline and language impairment in humans. A pathological hallmark is the accumulation of storage material within neuronal lysosomes resulting from mutations in the *CLN3* gene. We previously identified parallel deficits in auditory duration mismatch negativity (MMN), an electroencephalography (EEG)-based marker of auditory change detection, in individuals with CLN3 disease and in *Cln3*-/- mice. MMN-dependent auditory change detection relies on sensory-memory comparison mechanisms. However, the anatomical and neurophysiological substrates underlying this response in CLN3 disease remain unclear. Here, we investigated central auditory dysfunction in *Cln3*-/- mice by integrating immunohistochemical mapping of lysosomal storage pathology, using the canonical marker Subunit C of Mitochondrial ATP Synthase (SCMAS), with EEG analysis of auditory evoked potentials (AEPs). Neuropathological analyses revealed age-dependent and sex-divergent SCMAS accumulation across the auditory thalamocortical circuit, including the excitatory auditory thalamus, the inhibitory thalamic reticular nucleus, and the primary auditory cortex. In parallel, *Cln3*-/- mice exhibited age- and sex-dependent alterations in AEPs relative to wild-type controls. Importantly, an integrated measure of auditory thalamocortical SCMAS accumulation accounted for a substantial portion of age- and sex-matched variation in AEP responses, with stronger associations for the early N1 component than the later MMN component. Together, these findings link age-dependent and sex-divergent auditory neurophysiological deficits to region-specific lysosomal storage pathology within the auditory thalamocortical circuit in the *Cln3*-/- mouse model. This integrated functional-anatomical framework provides insight into circuit vulnerability and supports the development of translational neurophysiological biomarkers for CLN3 disease.

**Graphical Abstract:** 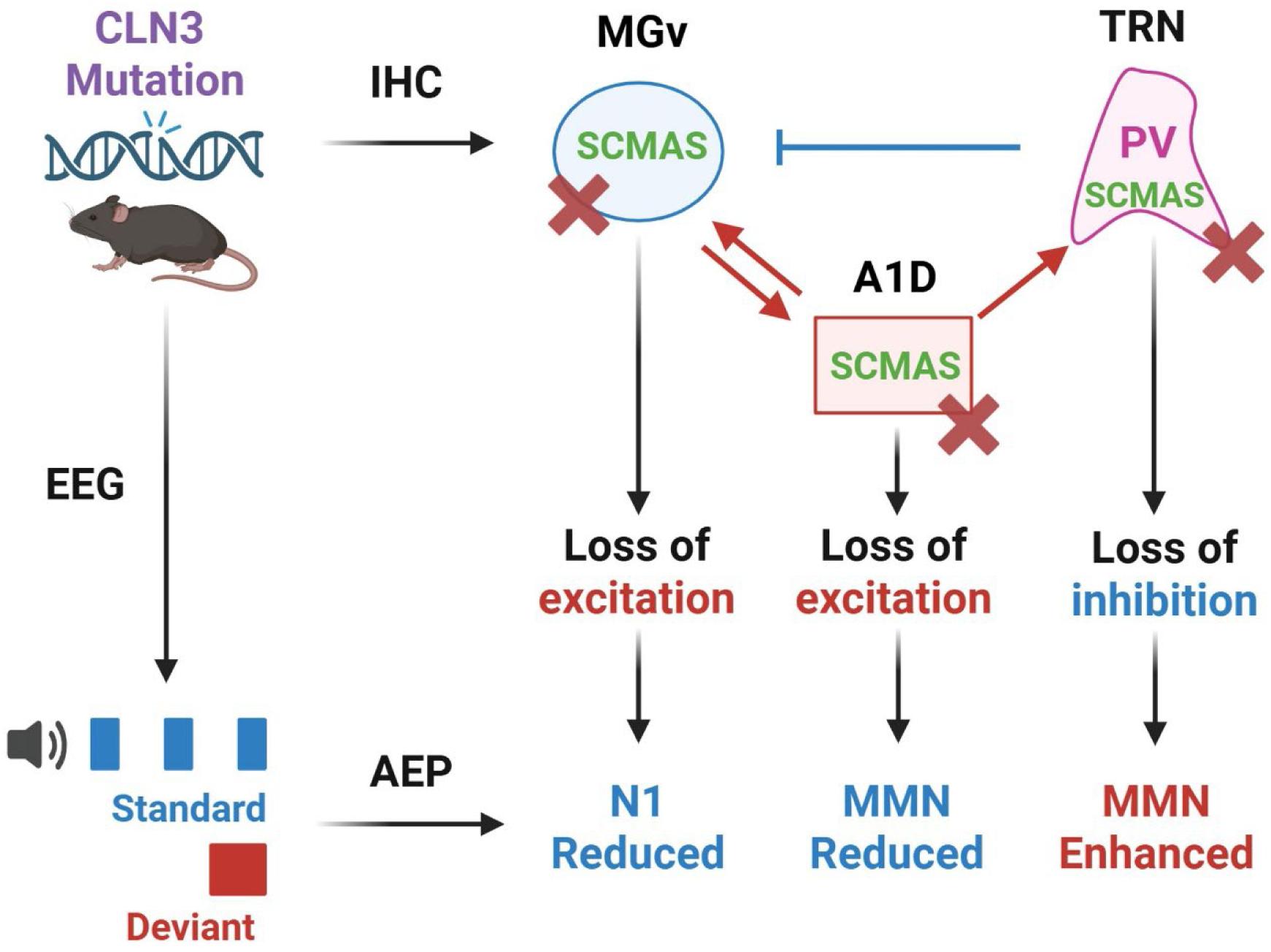

**Highlights:**

- SCMAS accumulates across the auditory thalamocortical circuit in *Cln3*-/- mice
- Accumulation shows distinct age- and sex-dependent regional trajectories
- *Cln3*-/- mice exhibit progressive alterations in auditory evoked potentials
- Thalamic pathology is strongly associated with the early auditory N1 response
- Histology–EEG integration links circuit pathology to auditory dysfunction

## 1. Introduction

Lysosomal Storage Disorders (LSDs) are inherited neurodegenerative diseases in which impaired lysosomal function leads to progressive accumulation of undegraded substrates and widespread neuronal dysfunction (Bellettato and Scarpa, 2010; Platt et al., 2018). Neuronal Ceroid Lipofuscinoses (NCLs), or Batten diseases, are a subset of LSDs marked by progressive accumulation of ceroid lipofuscin, particularly in neurons (Bennett and Hofmann, 1999; Mole et al., 1999; Johnson et al., 2019). CLN3 disease, caused by mutations in the *ceroid lipofuscinosis neuronal 3 (CLN3)* gene, is the juvenile-onset and most prevalent form of NCL (Consortium, 1995; Munroe et al., 1997; Kim et al., 2022). Clinical manifestations generally arise between 4 and 7 years of age with progressive vision loss (Adams et al., 2013; Ostergaard, 2016), followed by cognitive decline and language impairment, ultimately resulting in premature mortality around the second decade of life (Adams et al., 2007; Mink et al., 2013; Elmerskog et al., 2020; Tang et al., 2021; Morison et al., 2025). Sex-dependent differences in disease course have also been reported, whereby female patients tend to exhibit delayed clinical onset but faster progression (Cialone et al., 2012; Nielsen and Ostergaard, 2013). Because early visual impairment complicates standard cognitive assessments, objective neurophysiological biomarkers are critically needed to track disease progression.

The auditory system provides a promising avenue for such biomarkers. Auditory information ascends through a well-defined pathway, traveling from the cochlea through brainstem nuclei to the medial geniculate nucleus (MGN) of the thalamus. This signal is modulated by the inhibitory thalamic reticular nucleus (TRN) and ultimately arrives at the primary auditory cortex (A1) (Biacabe et al., 2001; Appler and Goodrich, 2011; Lee, 2013; Beebe et al., 2018; Suthakar and Liberman, 2021; Tomioka et al., 2023). Neural responses along this pathway can be readily measured noninvasively using electroencephalography (EEG), providing a translational window into auditory circuit function (Pruvost-Robieux et al., 2022). Auditory duration mismatch negativity (MMN) is a higher-order component of auditory evoked potentials (AEPs) elicited when deviant stimuli interrupt a repetitive sequence of standard sounds, reflecting auditory change detection (Ritter et al., 2002; Molholm et al., 2005; Naatanen et al., 2007). Our previous study in patients with CLN3 disease demonstrated clear reductions in the auditory duration MMN (Brima et al., 2024). A parallel study in *Cln3*-/- mice revealed similar deficits with sex- and age-dependent trajectories in MMN, while peripheral auditory function measured by auditory brainstem responses (ABR) remained intact (Ding et al., 2025). Together, these findings identify auditory MMN as a cross-species biomarker of central auditory dysfunction in CLN3 disease. However, the anatomical and cellular substrates linking lysosomal storage pathology to these neurophysiological deficits remain largely unknown.

Although lysosomal storage accumulation is a defining feature of CLN3 disease, its regional distribution within the auditory system has not been systematically characterized. Most previous neuropathological studies have focused on visual, motor, or cognitive pathways (Weimer et al., 2006; Ding et al., 2011; Burkovetskaya et al., 2017). One commonly used marker of CLN3 pathology is Subunit C of Mitochondrial ATP Synthase (SCMAS), a mitochondrial protein that accumulates within lysosomal storage deposits across multiple forms of NCL (Hall et al., 1991; Elleder et al., 1997; Gomez-Giro et al., 2019; Jana et al., 2023; Banach-Petrosky et al., 2025). While previous work has reported sex differences in SCMAS accumulation across several non-auditory brain regions at a single age point in a *Cln3* mouse model (Centa et al., 2023), it remains unknown whether sex-dependent patterns of SCMAS accumulation emerge across age within the auditory thalamocortical circuit.

Beyond regional distribution, cell-type vulnerability within auditory circuits also remains poorly understood. Parvalbumin-positive (PV+) inhibitory interneurons represent a candidate population that may be especially susceptible to lysosomal dysfunction due to their high metabolic demands, which arise from their rapid firing rate and their key roles in the temporal coordination of cortical and thalamic network activity (Hu et al., 2014; Ruden et al., 2021). In particular, PV+ neurons within the TRN provide powerful inhibitory control over thalamocortical relay neurons and are critical for shaping sensory signal transmission (Clemente-Perez et al., 2017; Steullet et al., 2018). Degeneration of cortical GABAergic interneurons, including PV+ cells, has been reported in *Cln3* mouse models at later disease stages (12-14 months of age) (Pontikis et al., 2005). However, lysosomal storage accumulation within these cells at earlier disease stages, prior to neurodegeneration, has not been systematically examined. Characterizing SCMAS accumulation within PV+ interneurons along the auditory thalamocortical circuit may therefore reveal early cellular vulnerabilities underlying auditory circuit pathology in *Cln3* mouse models.

Understanding how lysosomal storage pathology relates to circuit function is particularly important because different components of the AEP reflect distinct levels of auditory processing. The earliest AEP responses (0-10 ms) correspond to auditory brainstem activity, whereas the subsequent N1 component arises largely from thalamocortical transmission between MGN and A1 (Paiva et al., 2016; Modi and Sahin, 2017). Later components, such as MMN, reflect higher-order auditory integration across distributed cortical networks (Garrido et al., 2009; Lakatos et al., 2020). Disruptions of excitation–inhibition balance within the auditory thalamocortical circuit can therefore alter the amplitude and timing of specific AEP components (Light et al., 2010; Ross and Hamm, 2020; Zinnamon et al., 2023; Mazer et al., 2024; Du et al., 2025; Bayat et al., 2026). Although lysosomal dysfunction has been linked to mitochondrial impairment and synaptic abnormalities in CLN3 disease, it remains unresolved whether the emergence of neurophysiological deficits is directly associated with lysosomal storage accumulation (Ballabio and Gieselmann, 2009; Grunewald et al., 2017; Plotegher and Duchen, 2017; Stepien et al., 2020; Wang et al., 2026). Establishing this relationship would provide a mechanistic link between lysosomal storage pathology and circuit-level brain dysfunction, clarifying how cellular pathology within neural circuits manifests as measurable neurophysiological deficits.

In the present study, we combined histopathological mapping of lysosomal storage with longitudinal EEG measurements of auditory responses to determine how pathology within the auditory thalamocortical circuit relates to emerging neurophysiological dysfunction in *Cln3-/-* mice. SCMAS and PV immunostaining were used to quantify lysosomal storage and interneuron vulnerability in key auditory regions (TRN, MGN, and A1) across early stages of disease progression. In parallel, surface EEG recordings were used to measure AEPs longitudinally across the same age range. By integrating these approaches, we tested whether age- and sex-dependent patterns of lysosomal storage accumulation emerge across the auditory thalamocortical circuit and whether they are associated with alterations in auditory neurophysiology in *Cln3-/-* mice. This integrated analysis links lysosomal storage pathology within auditory circuits to emerging neurophysiological dysfunction, providing a foundation for translational EEG biomarkers in CLN3 disease.

## 2. Materials and Methods

### 2.1 Animals

All experimental procedures followed the National Institutes of Health (NIH) Guide for the Care and Use of Laboratory Animals and were approved by the University Committee on Animal Resources (UCAR) at the University of Rochester Medical Center (URMC). *Cln3-/-* mice and their wild-type (WT) littermates of both sexes were obtained from an in-house breeding colony derived from a line originally obtained from the Jackson Laboratory (B6.129S6-*Cln3*tm1Nbm/J, JAX:029471, ME). The *Cln3-/-* mouse strain was initially generated on the 129S6 genetic background (Mitchison et al., 1999) then back-crossed onto the C57BL/6J background for 10 generations before depositing at the Jackson Laboratory. Genotyping was performed by Transnetyx (Cordova, TN) using a real-time PCR-based assay, in accordance with the strain-specific protocol provided by the Jackson Laboratory. Animals were maintained under standard housing conditions with ad libitum access to food and water in a temperature- and humidity-controlled facility on a 12h:12h light–dark cycle. No overt signs of distress were noted in any animals during or following EEG recordings.

### 2.2 Immunohistochemistry (IHC)

#### 2.2.1 Perfusion and Sectioning

Animals were anesthetized with ketamine/xylazine (KX; K: 100 mg/kg; X: 10 mg/kg) and perfused with phosphate buffered saline (1X PBS) and 4% paraformaldehyde (PFA). Then, brains were dissected and fixed in 4% PFA for 24 hours before being stored in 1X PBS with 0.1% sodium azide (NaN_3_) at 4°C. Brain tissue was sectioned with a VF-310-0Z Compresstome (Precisionary Instruments, MA) in the coronal plane at a thickness of 80 μm. Slices were stored in 1X PBS with 0.1% NaN_3_ at 4°C in 24-well tissue culture plates (Thermo Fisher Scientific Inc., MA). Brain sections from *Cln3*-/- and WT mice of both sexes were collected at 3, 5, 7, and 9 months of age (n = 4 per age-, genotype- and sex-group).

#### 2.2.2 Immunostaining

The auditory TRN, MGv, and A1 slices were selected for immunostaining to assess central auditory neuropathology. The exact location of the auditory TRN was determined according to a prior study using retrogradely labeled neurons traced from different thalamic relay nuclei(Li et al., 2020). Immunostaining was performed on free-floating brain sections in 24-well plates using two to three slices per well. Slices were first washed in 1X PBS three times in 5 minutes. Non-specific protein binding was blocked using 5% normal goat serum (NGS) in 1X PBS with 0.01% Triton X-100 (PBS-T) for 1 hour. After blocking, slices were incubated overnight at 4°C in primary antibodies against SCMAS (1:250, anti-rabbit, ab181243, Abcam, MA) and PV (1:1000, anti-mouse, PV235, Swant, Switzerland) diluted in 5% NGS + PBS-T. Next, slices were washed 3 x 5 minutes in 1X PBS followed by 1 hour incubation in secondary antibodies (goat anti-rabbit Alexa Fluor 594: 1:500 and goat anti-mouse Alexa Fluor 647: 1:500, Thermo Fisher Scientific Inc., MA) diluted in 5% NGS + PBS-T. To ensure robust visualization of brain cells, slices were washed 3 x 5 minutes in 1X PBS and incubated for 5 minutes in DAPI nuclei counterstain solution (1:100 from 300 µM intermediate solution, D1306, Thermo Fisher Scientific Inc., MA, diluted in 1X PBS). Lastly, slices were washed 3 x 5 minutes in 1X PBS, mounted with media (Invitrogen^TM^ ProLong^TM^ Glass Antifade Mountant, P36982, Thermo Fisher Scientific Inc., MA) and cover-slipped on slides.

#### 2.2.3 Imaging

After immunostaining, brain sections were imaged by an Olympus VS120 Slide Scanner (Evident Scientific, PA) at Center for Musculoskeletal Research (CMSR) (URMC, NY) and a Nikon A1R HD – Pikachu Confocal Microscope (Nikon Corporation, Japan) at Center for Advanced Microscopy & Nanoscopy (CALMN) (URMC, NY). The slide scanner was used to efficiently survey storage material accumulation at lower magnification and resolution. Images were taken at 10X magnification for full brain slice scanning at blue (DAPI), green (autofluorescence), red (SCMAS) and far-red (PV) channels. The confocal microscope was utilized to assess SCMAS accumulation and co-localization of SCMAS with PV in specific areas at higher magnification and resolution. Confocal images were taken at 20X (0.4 µm per pixel) for the auditory TRN, MGv and A1 at blue (DAPI), green (Autofluorescence), red (SCMAS) and far-red (PV) channels.

#### 2.2.4 Image analysis

Image analysis was performed to quantify SCMAS accumulation and co-localization of SCMAS with PV using ImageJ (Fiji) and customized scripts in MATLAB (Mathworks, MA). For SCMAS quantification, images from the SCMAS channel were adjusted to subtract background, using adaptive local thresholding with Sauvola’s algorithm, and binarized after background subtraction. Global thresholding such as Otsu’s algorithm was not applied since SCMAS signals in WT mice were very sparse and dim, which would fail bimodal assumption. SCMAS+ area per cell (SCMAS+ area per cell = total SCMAS+ area above the threshold / total number of cells; cell number determined by DAPI) was calculated to represent SCMAS accumulation. Then, images from the SCMAS and PV channel, after background subtraction, were merged to calculate the percentage of co-localization (% of SCMAS+ cells that were also PV+ = PV+ SCMAS+ cells / SCMAS+ cells; % of PV+ cells that were also SCMAS+ = PV+ SCMAS+ cells / PV+ cells; cell number determined by DAPI). The color of SCMAS channel was reassigned to green in order to visualize co-localization with PV in magenta.

### 2.3 Electroencephalography (EEG)

#### 2.3.1 Surgical procedure

Surgical procedure for EEG electrode implantation was performed as described in our previous study (Ding et al., 2025). In brief, animals were anesthetized with isoflurane in a stereotaxic frame and implanted with a 32-channel surface EEG electrode array (H32, NeuroNexus Technologies Inc., MI) secured with dental cement. Postoperative care and analgesia followed the same protocol.

#### 2.3.2 Acoustic Stimulation

Following surgery and recovery, animals were head-fixed and placed on a rotating disc with a digital head stage connected. Acoustic stimulation was also identical to the prior study (Ding et al., 2025) except for the interstimulus interval (ISI). Broadband noise (standard: 50 ms; deviant: 100 ms; 80dB SPL) were presented in a pseudorandom sequence in blocks of 1000 trials, using the TDT system (RZ6 Multi-I/O processor, MF1 speaker, and Synapse software, Tucker-Davis Technologies, FL). 850 trials were standard stimuli, and 150 trials were deviant stimuli. In this study, ISI was increased from 400 ms to 800 and 1600 ms (supplementary material), with hardware parameters unchanged.

#### 2.3.3 Data Acquisition and Analysis

EEG data acquisition, artifact rejection, and analysis were performed following the previously established procedures(Ding et al., 2025). 11 male WT, 8 male *Cln3*-/-, 6 female WT, and 7 female *Cln3*-/- mice were recorded longitudinally from 3 to 9 months of age at 1600 and 800 ms ISI. Three 5 months old male WT mice and two 5 months old male *Cln3*-/- mice, used for pilot analysis, were included in addition to longitudinally recorded animals. There were 3 to 5 recording sessions for each animal per age point. Data was first averaged at individual animal level then grouped to compute genotype-, age-, and sex-differences. Group-averaged AEP waveforms at one representative electrode (Ch21) were plotted to visualize sex- and age-matched genotype difference and age-related difference in the same animals. The amplitude of N1 peak, which is the most negative deflection in the AEP waveform within 35 to 45 ms post stimuli onset, was also extracted at Ch21 to examine age- and genotype-related differences of primary auditory responses in sex-matched animals. In addition, spatiotemporal statistical analysis of the whole AEP waveform was performed to systematically characterize differences across electrode positions and time windows.

### 2.4 Statistical analysis

Statistical analyses of the EEG data were performed and plotted using customized scripts in MATLAB (Mathworks, MA) and GraphPad Prism 10.2.3 (GraphPad Software, CA). To explore spatiotemporal dynamics, two-sided unpaired t tests of AEP were performed between sex and age-matched WT and *Cln3*-/- mice across each electrode and time bin combination (electrode: Ch1-32, time: −50 to 350 ms in 10 ms bins). The p-values were corrected for multiple comparisons by controlling the False Discovery Rate (FDR) using MATLAB bioinformatics toolbox. Results were considered significant if their FDR-corrected p-values were less than 0.05. T-statistics, instead of p-values, were plotted across all the electrodes and time bins using MATLAB to show the direction of changes. Significant results were plotted in dark red and blue, and non-significant results were plotted in light red and blue. The same analysis was also performed for age-related AEP differences in WT and *Cln3*-/- mice of both sexes.

In addition, N1 peak amplitude from Ch21 was analyzed using a repeated measures two-way ANOVA, followed by Tukey’s multiple comparisons for sex- and age-matched WT and *Cln3*-/- mice. Significance was defined as p < 0.05. Effect size was reported as partial eta-squared (η²_p_)(Cohen, 1973, 1988; Lakens, 2013). A small effect has been defined as η² = 0.01, a medium effect as η² = 0.06, and a large effect as η² = 0.14(Cohen, 1988).

IHC Data was first averaged at individual animal level for both hemispheres then grouped to compute age, genotype and sex differences. A three-way ANOVA for each auditory processing brain region, followed by Tukey’s multiple comparisons for each age-, genotype- and sex- group was computed. Significance was defined as p < 0.05. Effect size was reported as partial eta-squared.

### 2.5 Partial least square regression

To investigate the relationship between histological and electrophysiological data in *Cln3*-/- mice, partial least square regression (PLS) was performed. Histological variables included SCMAS accumulation in the MGv, auditory TRN, A1 superficial (A1S), and A1 deep layers (A1D). Electrophysiological variables included both N1 peak amplitude and auditory duration MMN (Ding et al., 2025) to explore the potential anatomical correlates with early central auditory response and higher-order auditory function. Since the histological and electrophysiological data were collected from different batches of animals, group level summaries (mean, standard error of the mean (SEM), sample size) were computed for each age x sex combination. Then PLS regression was performed using the group means, including two components that are linear combinations of the partially correlated histological variables and account for more than 50% of the variance in the electrophysiological variables. Monte Carlo simulations (10,000 iterations) were performed by sampling perturbed histological means from multivariate normal distributions parameterized by within group covariance matrices and sampling electrophysiological means according to their SEM. PLS models were refit for each iteration, and coefficient stability was quantified using sign consistency and 95% confidence intervals (CI).

## 3. Results

### 3.1 Progressive lysosomal storage accumulation across the auditory thalamocortical circuit in *Cln3-/-* mice

To determine how lysosomal storage pathology develops across the auditory thalamocortical circuit, we quantified SCMAS accumulation in key thalamic relay, inhibitory gating, and cortical nodes of this pathway.

#### 3.1.1 Early sex difference in MGv lysosomal storage accumulation

Lysosomal storage accumulation was first examined in the ventral medial geniculate nucleus (MGv), the principal excitatory thalamic relay of the auditory circuit(Lee, 2013). WT mice of both sexes showed low levels of SCMAS accumulation from 3 to 9 months of age (Fig. 1), consistent with modest age-related increases of lysosomal storage observed in normal brain aging(Yang and Wang, 2021). In contrast, *Cln3*-/- mice exhibited markedly elevated and progressive SCMAS accumulation, beginning at higher levels and increasing at faster rates than WT controls (Fig. 1). Binarized images used for SCMAS quantification are included in the supplementary materials (Supplementary Fig. 1).

**Fig. 1.**
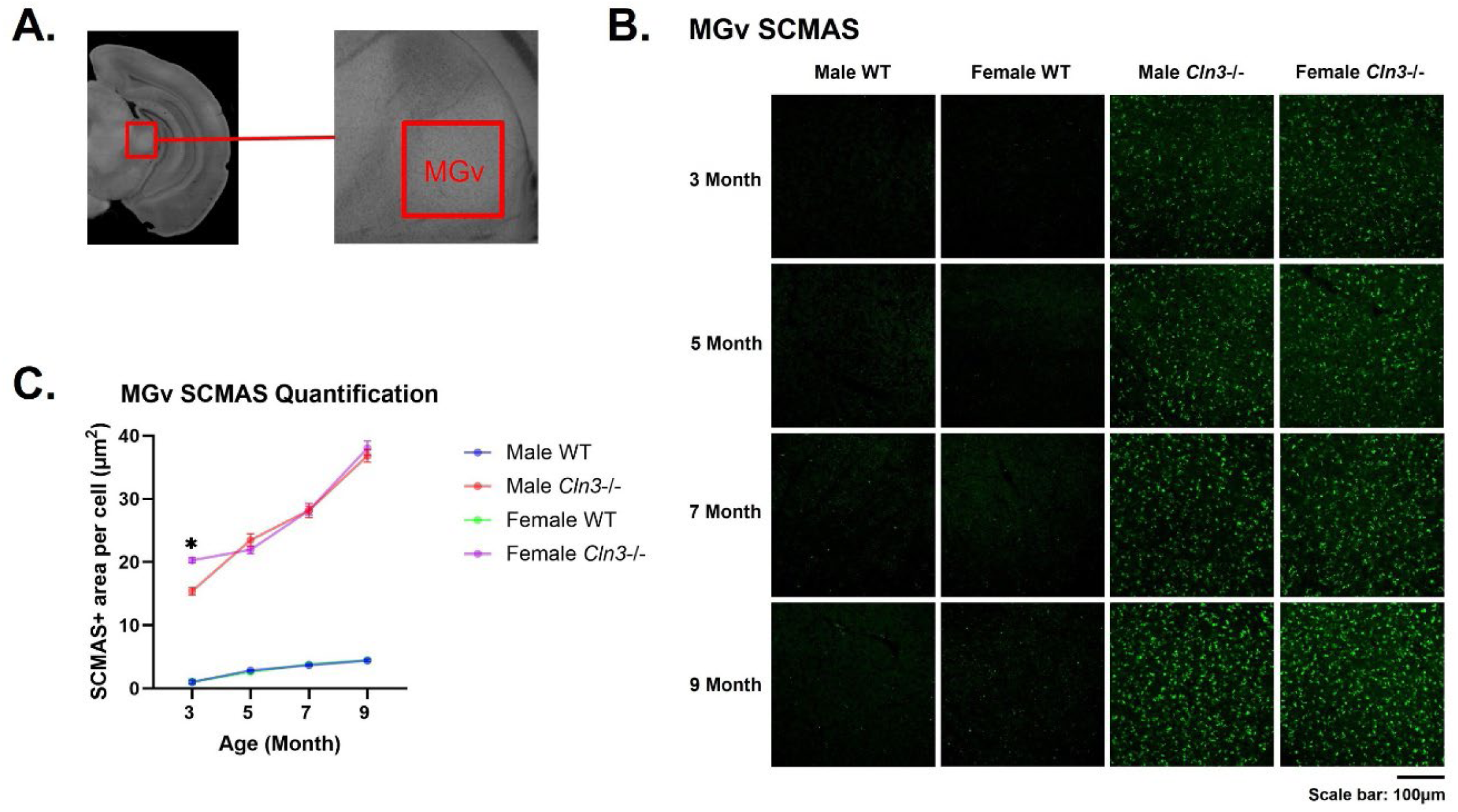
Early sex difference in MGv lysosomal storage accumulation. ***A,*** coronal brain section showing the location of the MGv. Red square indicates the image window. ***B,*** representative images of SCMAS accumulation (green) in male WT, female WT, male *Cln3*-/- and female *Cln3*-/- mice between 3 and 9 months of age. WT mice of both sexes showed minimal SCMAS accumulation. *Cln3*-/- mice of both sexes exhibited progressive SCMAS accumulation. Scale bar: 100μm. ***C,*** quantification of SCMAS+ area per cell in animals across age. Blue: male WT; Red: male *Cln3*-/-; Green: female WT; Purple: female male *Cln3*-/-. N = 4 mice per age, sex, and genotype group. Three-way ANOVA showed significant genotype (p < 0.0001), age (p < 0.0001), and genotype-by-age-by-sex interaction effect (p = 0.0073). Female *Cln3*-/- mice showed significantly higher SCMAS accumulation than male *Cln3*-/- mice at 3 months of age (p = 0.0001, Tukey’s test).

Three-way ANOVA showed significant effects of genotype (F(1,48) = 5556, p < 0.0001, η²_p_ = 0.991), age (F(3,48) = 241.1, p < 0.0001, η²_p_ = 0.938), genotype-by-age interaction ((F3,48) = 123.6, p < 0.0001, η²_p_ = 0.885), age-by-sex interaction (F(3,48) = 4.984, p = 0.0043, η²_p_ = 0.237), and genotype-by-age-by-sex interaction ((F3,48) = 4.505, p = 0.0073, η²_p_ = 0.220) (Fig. 1). Genotype accounted for the majority of variance, indicating robust pathological storage accumulation in *Cln3-/-* mice, with additional modulation by age and sex.

Post hoc comparisons revealed a significant sex difference at 3 months, with female *Cln3-/-* mice exhibiting higher SCMAS accumulation than males (p = 0.0001, Tukey’s multiple comparisons test) (Fig. 1). This sex difference disappeared at later ages (5, 7, and 9 months; all p > 0.9). Thus, female *Cln3-/-*mice show an early increase in lysosomal storage within the MGv that converges with males as pathology progresses.

#### 3.1.2 Late-emerging sex difference in lysosomal storage in the auditory TRN

SCMAS accumulation was next examined in the auditory TRN, an inhibitory gateway of thalamocortical transmission, receiving excitatory inputs from both MGv and deep layer auditory cortex and providing inhibitory outputs to the MGv(Pinault, 2004; Takata, 2020). The auditory TRN was defined using anatomical coordinates established in retrograde tracing studies of sensory thalamic nuclei(Li et al., 2020). Binarized images used for SCMAS quantification were included in supplementary materials (Supplementary Fig. 2).

While WT mice showed modest age-related increase of SCMAS, *Cln3*-/-mice displayed robust and progressive pathological SCMAS accumulation across age (Fig 2). Three-way ANOVA showed significant effects of genotype (F(1,48) = 19149, p < 0.0001, η²_p_ = 0.998), age (F(3,48) = 501.4, p < 0.0001, η²_p_ = 0.969), sex (F(1,48) = 13.55, p = 0.0006, η²_p_ = 0.220), genotype-by-age interaction (F3,48) = 212.9, p < 0.0001, η²_p_ = 0.930), age-by-sex interaction (F(3,48) = 22.45, p < 0.0001, η²_p_ = 0.584), sex-by-genotype interaction (F1,48) = 14.33, p = 0.0004, η²_p_ = 0.230), and genotype-by-age-by-sex interaction (F3,48) = 19.35, p < 0.0001, η²_p_ = 0.547) (Fig 2). Genotype again accounted for the largest proportion of variance, with modulation by age and sex.

**Fig. 2.**
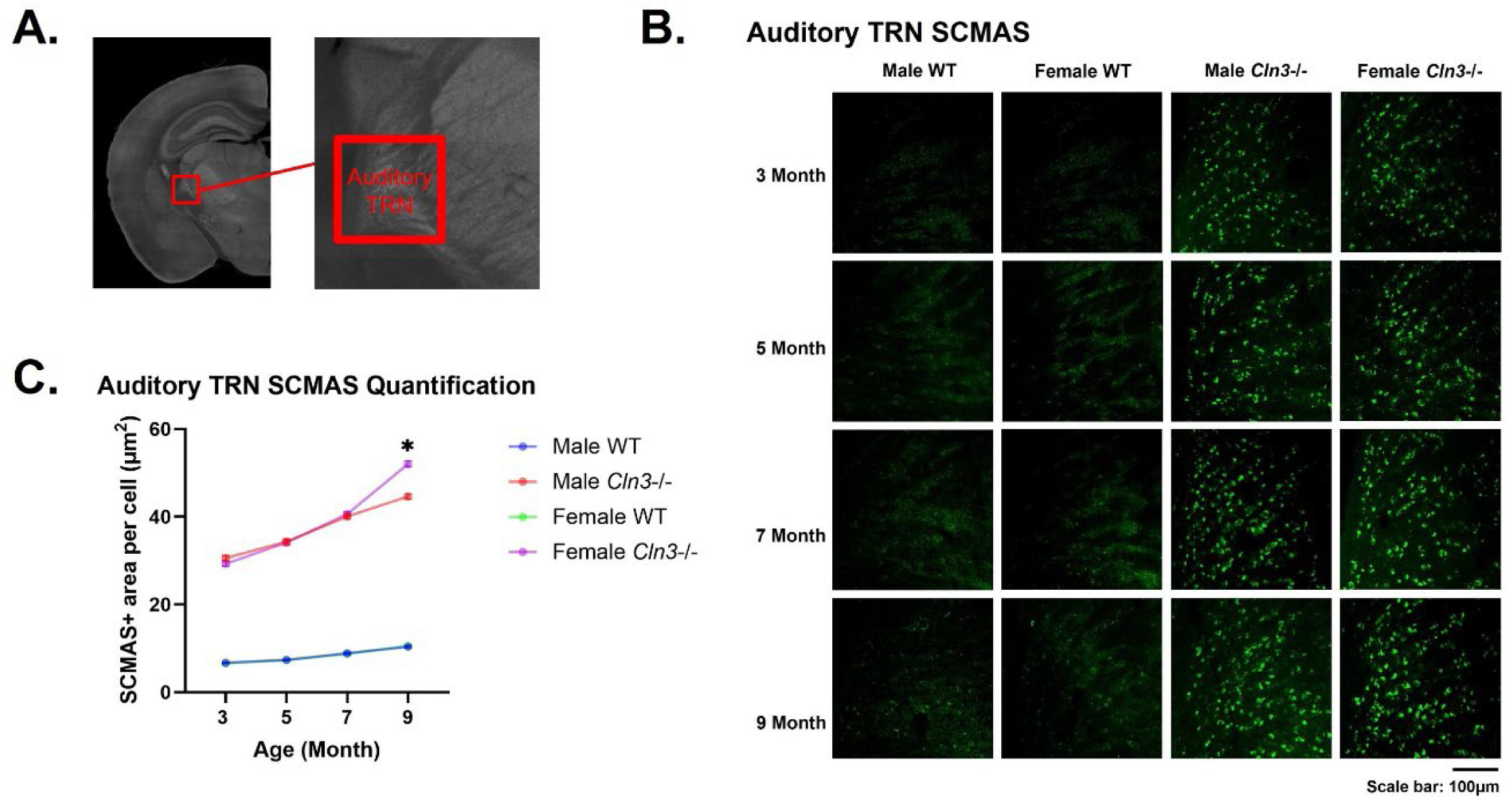
Late-emerging sex difference in lysosomal storage in the auditory TRN. ***A,*** coronal brain section showing the location of the auditory TRN. Red square indicates the image window. ***B,*** representative images of SCMAS accumulation (green) in male WT, female WT, male *Cln3*-/- and female *Cln3*-/- mice between 3 and 9 months of age. WT mice of both sexes showed minimal SCMAS accumulation. *Cln3*-/- mice of both sexes exhibited progressive SCMAS accumulation. Scale bar: 100μm. ***C,*** quantification of SCMAS+ area per cell in animals across age. Blue: male WT; Red: male *Cln3*-/-; Green: female WT; Purple: female male *Cln3*-/-. N = 4 mice per age, sex, and genotype group. Three-way ANOVA showed significant genotype (p < 0.0001), age (p < 0.0001), sex (p = 0.0006) and genotype-by-age-by-sex interaction effect (p < 0.0001). Female *Cln3*-/- mice showed significantly higher SCMAS accumulation than male *Cln3*-/- mice at 9 months of age (p < 0.0001, Tukey’s test).

No sex differences were observed in Cln3-/- mice at 3, 5, or 7 months (all p > 0.7, Tukey’s test). However, by 9 months female Cln3-/- mice exhibited significantly greater SCMAS accumulation than males (p < 0.0001; Tukey’s test) (Fig. 2). These results indicate a late-emerging sex difference in pathological lysosomal storage within the inhibitory TRN.

#### 3.1.3 Cortical lysosomal storage accumulation with dynamic sex differences in A1 deep layers

Next, SCMAS accumulation was assessed in A1, the primary cortical target of the auditory thalamus. Because superficial (L1-L4) and deep (L5-L6) cortical layers have distinct connectivity and functional roles in thalamocortical processing, from higher-order integration to feedback regulation (Smith et al., 2012; Slater and Isaacson, 2020), they were analyzed separately.

In superficial layers (L1–L4), *Cln3-/-* mice of both sexes showed progressive SCMAS accumulation between 3 and 9 months relative to WT controls (Fig. 3, Supplementary Fig. 3). Three-way ANOVA showed significant effects of genotype (F(1,48) = 10483, p < 0.0001, η²_p_ = 0.995), age (F(3,48) = 830.5, p < 0.0001, η²_p_ = 0.981), age-by-genotype interaction (F3,48) = 421.1, p < 0.0001, η²_p_ = 0.963), and sex-by-genotype interaction (F1,48) = 4.928, p = 0.0312, η²_p_ = 0.093). However, no significant sex effect, age-by-sex interaction, or age-by-sex-by-genotype interaction was observed.

**Fig. 3.**
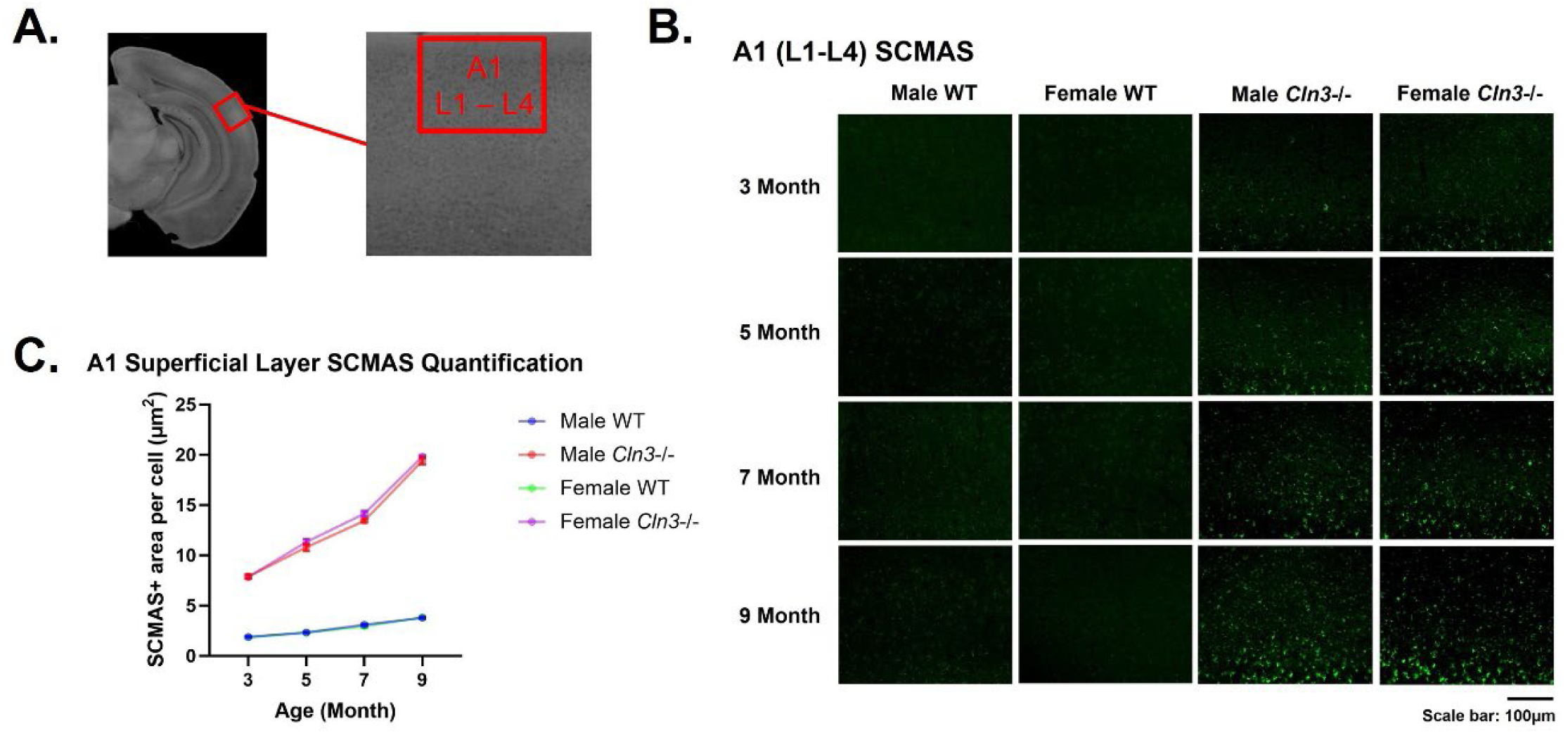
Lysosomal storage accumulation in A1 superficial layers is primarily driven by genotype and age rather than sex. ***A,*** coronal brain section showing the location of the A1 superficial layers (L1 – L4). Red square indicates the image window. ***B,*** representative images of SCMAS accumulation (green) in male WT, female WT, male *Cln3*-/- and female *Cln3*-/- mice between 3 and 9 months of age. WT mice of both sexes showed no pathological SCMAS accumulation. *Cln3*-/-mice of both sexes exhibited progressive SCMAS accumulation. Scale bar: 100μm. ***C,*** quantification of SCMAS+ area per cell in animals across age. Blue: male WT; Red: male *Cln3*-/-; Green: female WT; Purple: female male *Cln3*-/-. N = 4 mice per age, sex, and genotype group. Three-way ANOVA showed significant genotype (p < 0.0001) and age (p < 0.0001) effects. Sex effect was not significant (p = 0.0757). Male and female *Cln3*-/- mice showed no sex-specific difference in SCMAS accumulation in the A1 superficial layer.

Post hoc comparisons confirmed no sex differences in A1 superficial-layer SCMAS accumulation at any age (all p > 0.5, Tukey’s test). Thus, lysosomal storage accumulation in A1 superficial layers is primarily driven by genotype and age rather than sex.

In contrast to superficial layers, deep layers of A1 (L5–L6) exhibited both age- and sex-dependent patterns of SCMAS accumulation (Fig. 4, Supplementary Fig. 4). Three-way ANOVA showed significant effects of genotype (F(1,48) = 6467, p < 0.0001, η²_p_ = 0.993), age (F(3,48) = 496.8, p < 0.0001, η²_p_ = 0.969), sex (F(1,48) = 5.647, p = 0.0215, η²_p_ = 0.105), age-by-genotype interaction (F3,48) = 297.5, p < 0.0001, η²_p_ = 0.949), age-by-sex interaction (F(3,48) = 16.75, p < 0.0001, η²_p_ = 0.512), sex-by-genotype interaction (F1,48) = 5.722, p = 0.0207, η²_p_ = 0.107), and age-by-sex-by-genotype interaction (F3,48) = 17.9, p < 0.0001, η²_p_ = 0.528) (Fig. 4). Genotype remained the dominant contributor to variance, with significant modulations by age and sex.

**Fig. 4.**
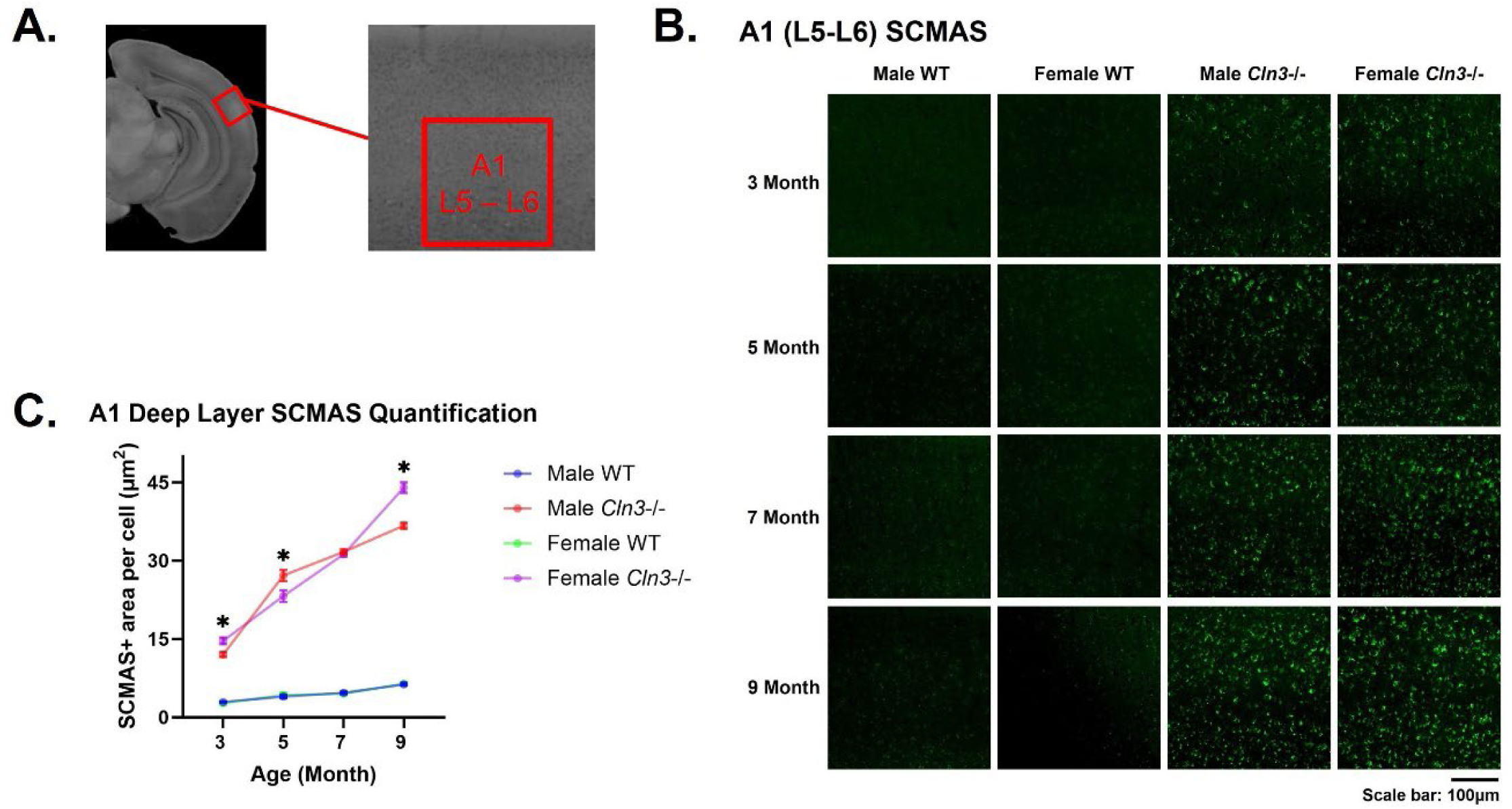
Dynamic and sex-dependent trajectory of lysosomal storage accumulation in A1 deep layers. ***A,*** coronal brain section showing the location of the A1 deep layers (L5 – L6). Red square indicates the image window. ***B,*** representative images of SCMAS accumulation (green) in male WT, female WT, male *Cln3*-/- and female *Cln3*-/- mice between 3 and 9 months of age. WT mice of both sexes showed minimal SCMAS accumulation. *Cln3*-/- mice of both sexes exhibited progressive SCMAS accumulation. Scale bar: 100μm. ***C,*** quantification of SCMAS+ area per cell in animals across age. Blue: male WT; Red: male *Cln3*-/-; Green: female WT; Purple: female male *Cln3*-/-. N = 4 mice per age, sex, and genotype group. Three-way ANOVA showed significant genotype (p < 0.0001), age (p < 0.0001), sex (p = 0.0215), and age-by-sex-by-genotype interaction effect (p < 0.0001). Male and female *Cln3*-/- mice showed sex-specific differences in SCMAS accumulation in the A1 deep layer at 3, 5 and 9 months of age (3 month: p = 0.0315; 5 month: p = 0.0012, 7 month: p > 0.9999; 9 month: p < 0.0001). Female *Cln3*-/-mice showed higher SCMAS at 3 and 9 months of age, while male *Cln3*-/- mice showed higher SCMAS at 5 months of age.

Post hoc analyses showed that female *Cln3-/-* mice exhibited higher SCMAS levels at 3 and 9 months (3 months: p = 0.0315; 9 months: p < 0.0001, Tukey’s test). Conversely, males showed a transient increase at 5 months (p = 0.0012, Tukey’s test), while no sex difference was detected at 7 months. These findings indicate a dynamic and sex-dependent trajectory of lysosomal storage accumulation in A1 deep layers.

Across ages, SCMAS accumulation was consistently greater in deep layers than in superficial layers, suggesting heightened vulnerability of deep-layer cortical neurons to lysosomal storage pathology. Because neurons in layers 5 and 6 provide corticofugal and corticothalamic projections that regulate thalamocortical sensory processing (Doucet et al., 2002; Meltzer and Ryugo, 2006; Znamenskiy and Zador, 2013; Homma and Bajo, 2021), preferential pathology in these layers may disrupt feedback control within the auditory thalamocortical circuit and contribute to abnormal central auditory processing.

Together, these results reveal region- and layer-specific patterns of lysosomal storage accumulation across the auditory thalamocortical circuit. Sex-divergences emerges early in the thalamic relay nucleus (MGv), later in the inhibitory TRN, and becomes particularly pronounced in deep cortical layers that provide corticothalamic feedback.

### 3.2 Cell type susceptibility to SCMAS accumulation in the *Cln3*-/- mouse model

To determine whether specific neuronal populations exhibit differential susceptibility to lysosomal storage pathology, we examined SCMAS accumulation in defined cell types across auditory thalamocortical regions in *Cln3-/-* mice.

In the auditory TRN, PV immunoreactivity densely co-localized with SCMAS in male and female *Cln3*-/- mice between 3 and 9 months of age (Fig. 5A). PV+ cells constituted the majority of DAPI+ cells in the auditory TRN, consistent with previous anatomical studies (Li et al., 2020; Luo et al., 2024). Around 90% of PV+ cells contained SCMAS+ clusters (# of SCMAS+ PV+ cells / # of PV+ cells) (Fig. 5C), indicating that auditory TRN PV+ interneurons represent a major cell type susceptible to pathological lysosomal storage accumulation. The percentage of SCMAS+ cells that were also PV+ in TRN (# of SCMAS+ PV+ cells / # of SCMAS+ cells) was about 80% in both male and female *Cln3*-/- mice across age (Fig. 5E).

**Fig. 5.**
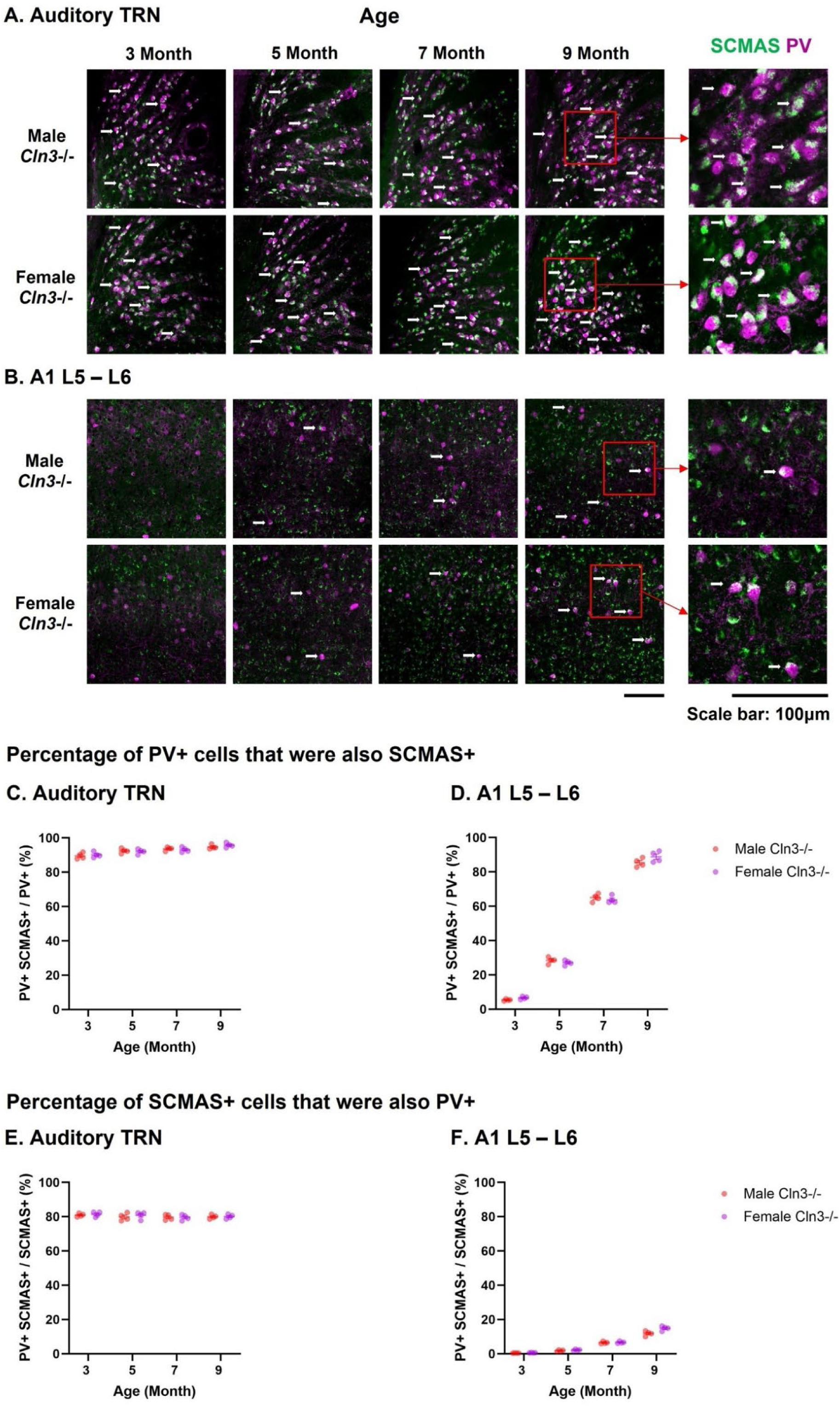
Cell type susceptibility to SCMAS accumulation in the *Cln3*-/- mouse model. ***A,*** PV and SCMAS co-localization in the auditory TRN in male and female *Cln3*-/- mice between 3 and 9 months of age. Magenta: PV; Green: SCMAS. White arrow: co-localization. The fifth column shows zoom-in images of PV and SCMAS co-localization in 9-month-old male and female *Cln3*-/- mice. Red square and arrow show the image windows for zoom-in. Scale bar: 100μm. ***B,*** PV and SCMAS co-localization in the A1 deep layer in male and female *Cln3*-/- mice between 3 to 9 months of age. ***C and D,*** percentage of PV+ cells that were also SCMAS+ (# PV+ SCMAS+ cells / # PV+ cells) across age. C: TRN; D: A1 deep layers. Red: male *Cln3*-/- mice. Purple: female *Cln3*-/- mice. ***E and F,*** percentage of SCMAS+ cells that were also PV+ (# PV+ SCMAS+ cells / # SCMAS+ cells) across age. E: TRN; F: A1 deep layers. Red: male *Cln3*-/- mice. Purple: female *Cln3*-/- mice.

Despite their relatively low abundance, PV+ cells in A1 also exhibited SCMAS accumulation (Fig. 5B). Co-localization analysis focused on the A1 deep layers, where *Cln3*-/- mice showed more pronounced SCMAS accumulation than superficial layers (Figs. 3, 4). PV+ cells constituted a minority of cells in L5-L6 of A1 (Fig. 5B), consistent with previous reports (Yuan et al., 2011). At 3 months of age, there was minimal co-localization of PV and SCMAS in A1 of *Cln3*-/- mice (Fig. 5B, D, F). Instead, SCMAS+ clusters were predominantly observed in PV-negative neurons, which likely correspond to excitatory neurons that constitute the majority (>80%) of cortical neurons (Wang et al., 2018). Between 5 and 9 months of age, the percentage of PV+ cells that were SCMAS+ progressively increased in L5-L6 of A1 in *Cln3*-/- mice (Fig. 5D). Similarly, the percentage of SCMAS+ cells that were PV+ showed a gradual increase over time, although it remained low overall (Fig. 5F). Together, these results suggest that the sex-dependent differences in deep-layer A1 pathology arise primarily from PV- neuronal populations.

In rodents, the MGv is mainly composed of glutamatergic relay neurons that provide excitatory thalamocortical input to auditory cortex (Tomioka et al., 2023). Consistent with this cellular composition, no PV+ cell or PV-SCMAS co-localization were observed in the MGv (Supplementary Fig. 5). Instead, SCMAS+ clusters were present in excitatory neurons.

Together, these results identify cell-type–specific susceptibility to lysosomal storage pathology within the auditory circuit: PV+ inhibitory interneurons in the TRN and excitatory neurons in both MGv and deep-layer A1 show prominent SCMAS accumulation in *Cln3-/-* mice. Accumulation within these populations may disrupt excitation–inhibition balance within the auditory thalamocortical circuit.

### 3.3 WT mice exhibit stable auditory evoked potentials across age

Before examining lysosomal storage pathology-related neurophysiological changes, we first assessed whether auditory evoked potentials (AEPs) could be reliably recorded across age in WT mice. Animals were implanted with surface EEG arrays to record AEP longitudinally across the same age range (3-9 months) (Fig. 5A, B). Acoustic stimulation was similar to our previous MMN study(Ding et al., 2025), including both standard stimuli (50 ms, 85% trials) and deviant stimuli (100 ms, 15% trials). However, the interstimulus interval (ISI) was expanded from 400 ms in the MMN study to 800 ms and 1600 ms, reducing response adaptation during repeated stimulus presentation and enabling more robust assessment of the N1 component that primarily reflects thalamocortical transmission (Gonsalvez and Polich, 2002; Vallesi et al., 2013).

Focusing on a centrally located electrode (Ch21), both male and female WT mice displayed robust and stable AEP responses to standard stimuli at 1600 ms ISI from 3 to 9 months of age (Fig. 6C, D). Standard AEP waveforms showed minimal age-related variation. Similarly, deviant AEP responses showed little age-dependent change in both sexes (Supplementary Fig. 6). The amplitude of standard and deviant AEP responses is comparable, suggesting no detection of duration change over this long ISI.

**Fig. 6.**
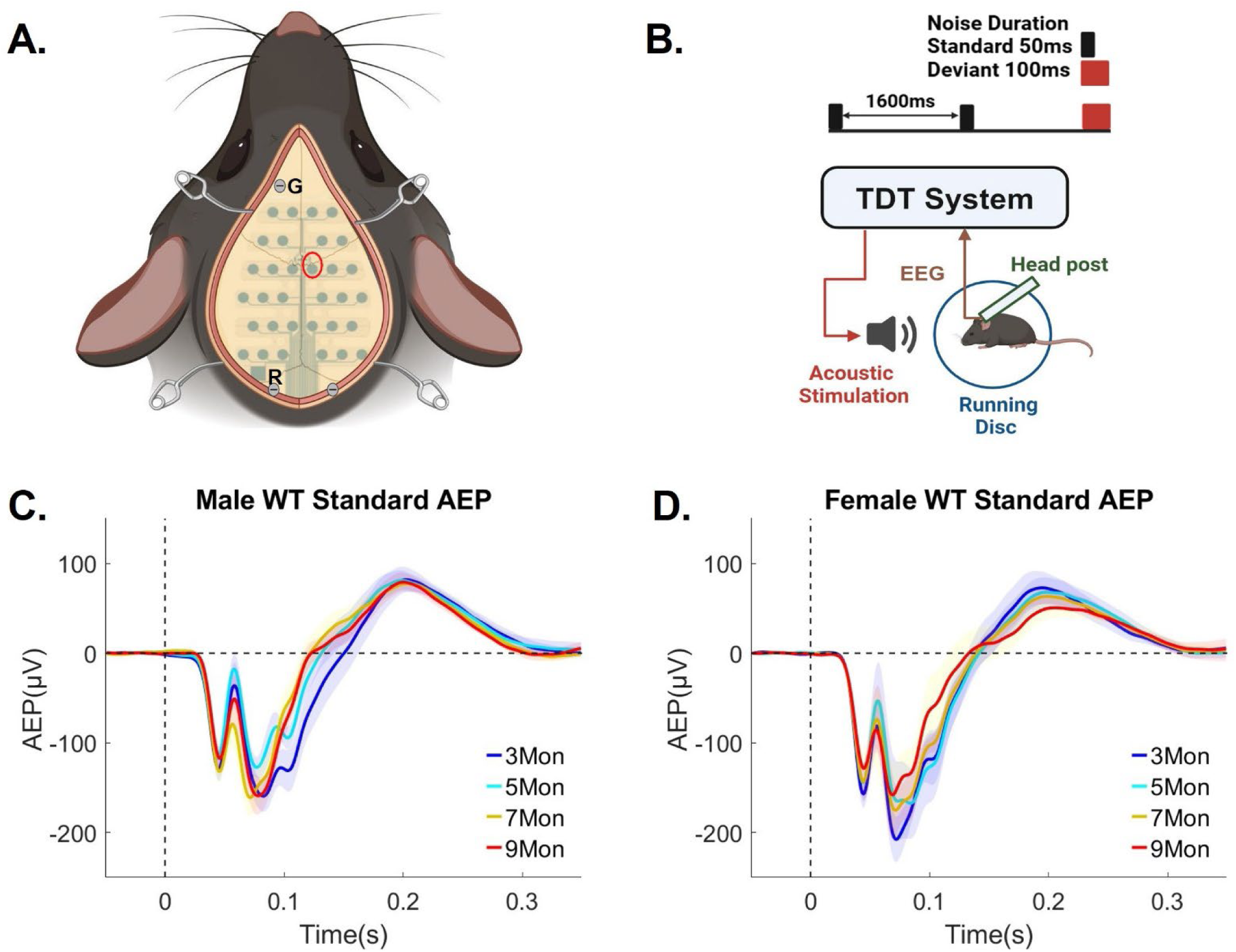
WT mice of both sexes showed consistent AEP across age. ***A,*** Diagram of the 32- channel mouse EEG electrode array implanted on the skull of a mouse. The cross symbol on the array is positioned at the bregma. Screws are inserted for grounding (“G”), reference (“R”), and probe anchorage. Red circle shows the location of Ch21 (representative auditory duration MMN and AEP waveforms taken from this channel). ***B***, Diagram shows the setup for recording AEP in mice. Animals are head-fixed to a head-post, with EEG and locomotion data recorded using a TDT system. A speaker is placed 10 cm in front of the animal. Standard tone: 50 ms duration, 850 trials/session; Deviant tone: 100 ms duration, 150 trials/session. Interstimulus interval (ISI): 1600 ms. ***C and D*,** Trial and subject averaged standard AEP waveforms for male (C) and female (D) WT mice aged 3 to 9 months, recorded from a centrally located channel (Ch21). The vertical dashed line indicates stimulus onset. AEPs in response to the standard tone are presented, with the shaded areas representing the standard error of the mean (SEM). Blue: 3 month; Cyan: 5 month, Yellow: 7 month, Red: 9 month. Male WT: n = 11 mice for 3-, 7- and 9- month-old groups; n = 14 for the 5-month old group. Female WT: n = 6 mice for all ages. All animals were recorded longitudinally from 3 to 9 months of age, except three additional male mice recorded only at 5 months of age in pilot studies. Male and female WT mice showed robust AEP with no significant age-related changes.

Analysis at the 800 ms ISI produced similar results, indicating that WT mice of both sexes exhibited consistent AEP responses across age (Supplementary Fig. 7). Compared to the 1600 ms ISI, AEP waveforms at the 800 ms ISI showed evidence of repetition suppression, in which the negativity of the auditory response decreased as the interval between stimuli shortened (Vallesi et al., 2013; Fitzgerald and Todd, 2020).

Overall, WT mice exhibited stable auditory neurophysiological responses across age, supporting the reliability of the EEG recording paradigm for longitudinal assessment of auditory function.

### 3.4 *Cln3*-/- mice show sex-specific and age-dependent auditory neurophysiological deficits

#### 3.4.1 Male *Cln3-/-* mice exhibit progressive reduction of the N1 from 5-9 months

Next, age-related changes of auditory neurophysiological response were examined in the *Cln3*-/- mouse model. Compared to male WT mice, male *Cln3*-/-mice showed no difference in N1 amplitude at 3 months, followed by a progressive reduction between 5 and 9 months of age (Fig. 7).

**Fig. 7.**
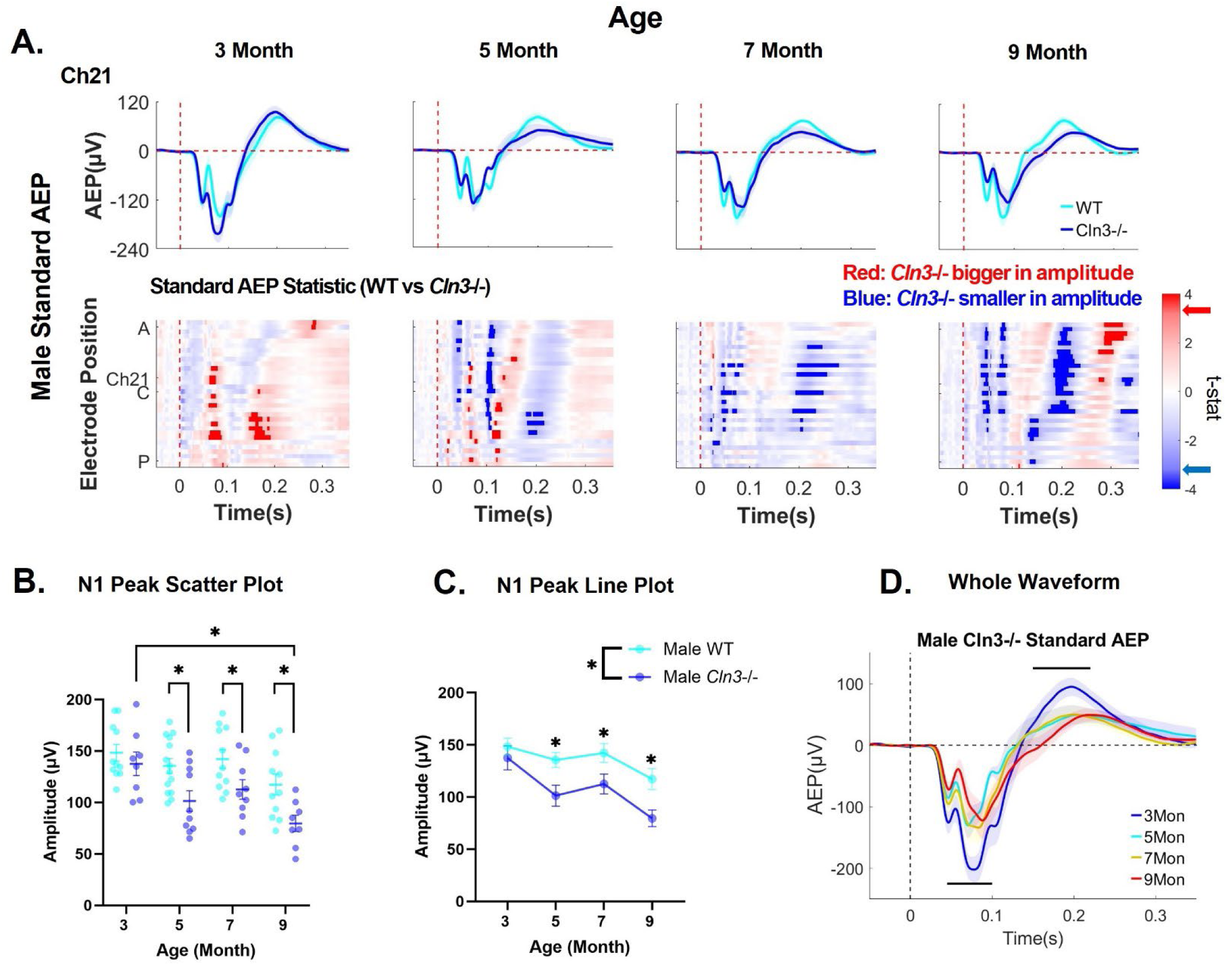
Male *Cln3*-/- mice exhibit progressive reduction of the N1 from 5-9 months. ***A*,** top row, trial- and subject-averaged standard AEP waveforms for male WT mice (light blue) and male *Cln3*-/- mice (dark blue) at Ch21. The vertical dashed red line indicates stimulus onset. Bottom row, statistical differences in standard AEP between male WT and *Cln3*-/- mice were displayed across all electrodes and the entire trial duration (32 electrodes × 400ms). Results from t-tests were corrected for multiple comparisons using the FDR method. Significance was determined when the FDR-adjusted p-value was less than 0.05 (two-tailed). Electrode positions over the mouse skull are indicated from A (anterior) to C (center) and P (posterior). Red indicates electrodes and time bins (in 20ms) where *Cln3*-/-mice showed larger amplitudes (further away from zero compared to WT mice), while blue indicates electrodes and time bins where *Cln3*-/- mice showed smaller amplitudes (closer to zero compared to WT mice). Significant t-stat values are represented by dark red and dark blue, based on the FDR-corrected p-value. Red and blue arrows on the color bar indicate significant t-stat value. Male WT: n = 11 mice for 3-, 7- and 9-month-old groups; n = 14 for the 5-month-old group. Male *Cln3*-/-: n = 8 mice for 3-, 7- and 9-month-old groups; n = 10 for the 5-month- old group. ***B and C,*** scatter and line plots of N1 amplitude (Ch21) for male WT and *Cln3*-/- mice. Individual data points are for each animal and error bars represent mean ± SEM. Repeated measures two-way ANOVA showed a significant genotype effect (p < 0.0001), with a progressive reduction in N1 in male *Cln3*-/- mice compared to male WT mice. Age-matched genotype comparison showed that male *Cln3*-/- mice exhibited reduction in N1 at 5, 7 and 9 months of age (3 month: p = 0.4273, 5 month: p = 0.0059, 7 month: p = 0.0266, 9 month: p = 0.0069). ***D,*** Trial and subject averaged standard AEP waveforms for male *Cln3*-/- mice from 3 to 9 months of age (Ch21). The vertical dashed line indicates stimulus onset. Shaded areas represent the standard error of the mean (SEM). Blue: 3 month; Cyan: 5 month, Yellow: 7 month, Red: 9 month. The black horizontal lines indicate the time windows during which AEP at 9 months is significantly reduced compared to AEP at 3 months (p < 0.05, FDR corrected unpaired t-tests).

Spatiotemporal analysis revealed that 3-month-old male *Cln3*-/- mice displayed no significant difference in standard AEP around the N1 (∼40 ms) (p > 0.05, FDR-corrected unpaired t-tests) (Fig. 7A). However, increased activity was observed around 70-90 ms and 160-190 ms at central-posterior electrodes (p < 0.05, FDR-corrected unpaired t-tests).

Starting at 5 months, male *Cln3*-/- mice showed significant reductions in AEP amplitude around the N1 (∼40 ms) and late positive peak (∼200 ms) (p < 0.05, FDR-corrected unpaired t-tests) (Fig. 7A). These deficits were most pronounced at 9 months, particularly at anterior-central electrodes around 40, 80 and 200 ms post stimulus onset (p < 0.05, FDR-corrected unpaired t-tests).

N1 peak analysis confirmed that male *Cln3*-/- mice displayed a progressive reduction in N1 amplitude (Fig. 7B and 7C). Two-way ANOVA revealed a significant genotype effect (F(1, 74) = 18.43, p < 0.0001, η²_p_ = 0.217) (Fig. 7B and 7C). Compared to age-matched male WT mice, male *Cln3*-/- mice exhibited no difference at 3 months (p = 0.4273, Tukey’s test) but significant decrease at 5, 7, and 9 months of age (5 month: p = 0.0059; 7 month: p = 0.0266; 9 month: p = 0.0069, Tukey’s test).

Age also had a significant effect across pooled genotypes (F(3, 74) = 7.536, p = 0.0002, η²_p_ = 0.235, two-way ANOVA) (Fig. 7B and 7C). While male WT mice showed no significant age-related decline (p > 0.05, Tukey’s test), male *Cln3*-/-mice exhibited significant N1 reduction at 9 months relative to 3 months (p = 0.0009, Tukey’s test) (Fig. 7B and 7C). Whole AEP waveform analysis further confirmed age-dependent decline of AEP responses, with significant reductions between 40–100 ms and 150–220 ms at 9 months compared with 3 months (p < 0.05, FDR-corrected unpaired t-tests) (Fig. 7D).

Additional analyses of AEP responses to deviant stimuli (Supplementary Fig. 8) and responses at 800 ms ISI (Supplementary Fig. 9) revealed consistent genotype- and age-dependent differences. Together, these results demonstrate progressive reduction of the N1 component in male *Cln3-/-* mice beginning at 5 months of age.

#### 3.4.2 Female *Cln3*-/- mice exhibit early-onset and progressive reduction of the N1

Next, N1 alterations in female *Cln3-/-* mice were investigated. Compared to female WT mice, female *Cln3*-/- mice showed a progressive reduction in N1 amplitude beginning at 3 months of age (Fig. 8).

**Fig. 8.**
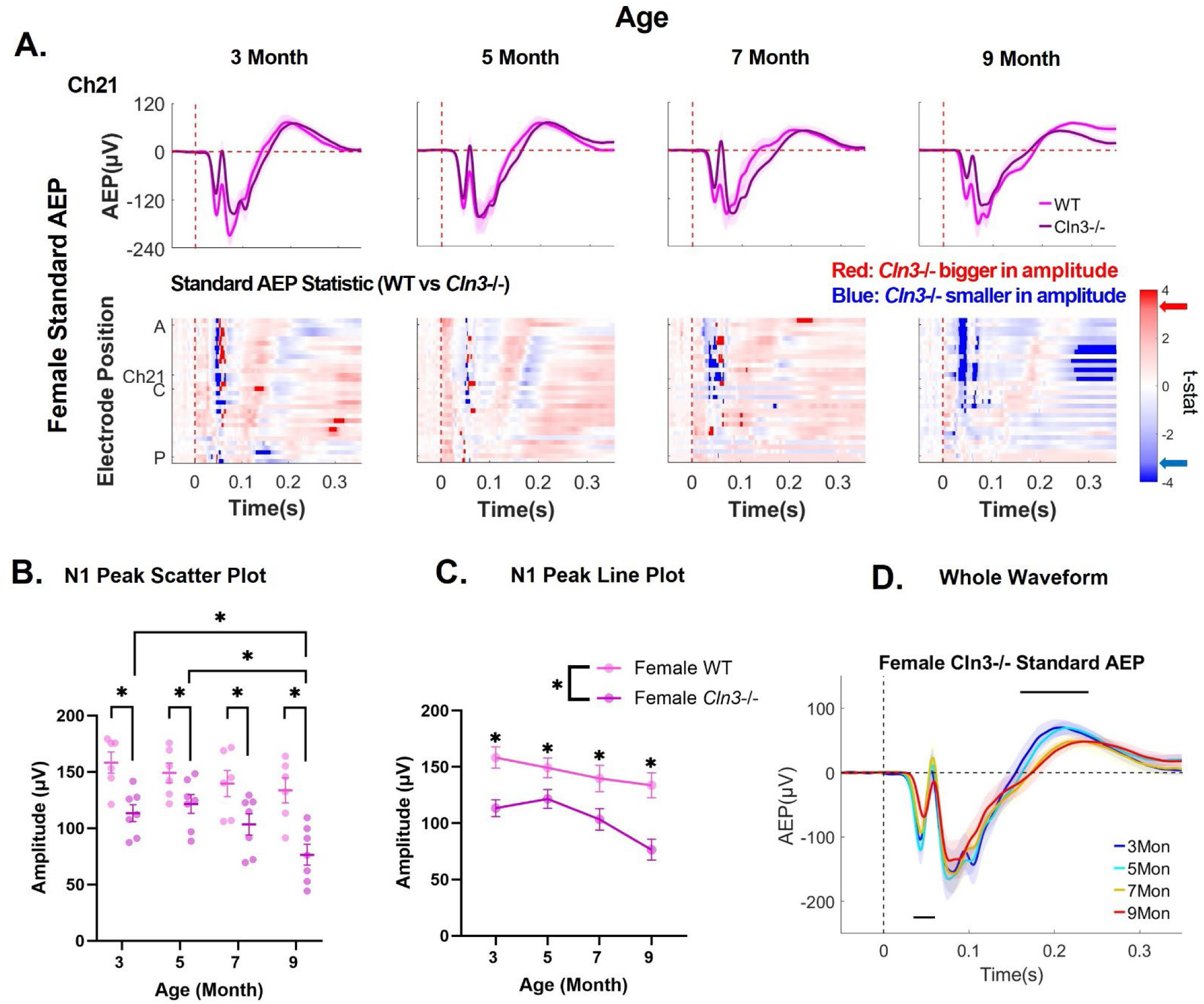
Female *Cln3*-/- mice exhibit early-onset and progressive reduction of the N1. ***A*,** top row, trial- and subject-averaged standard AEP waveforms for female WT mice (light pink) and female *Cln3*-/- mice (dark pink) at Ch21. The vertical dashed red line indicates stimulus onset. Bottom row, statistical differences in standard AEP between female WT and *Cln3*-/- mice were displayed across all electrodes and the entire trial duration (32 electrodes × 400ms). Results from t-tests were corrected for multiple comparisons using the FDR method. Significance was determined when the FDR-adjusted p-value was less than 0.05 (two-tailed). Electrode positions over the mouse skull are indicated from A (anterior) to C (center) and P (posterior). Red indicates electrodes and time bins (in 20ms) where *Cln3*-/-mice showed larger amplitudes (further away from zero compared to WT mice), while blue indicates electrodes and time bins where *Cln3*-/- mice showed smaller amplitudes (closer to zero compared to WT mice). Significant t-stat values are represented by dark red and dark blue, based on the FDR-corrected p-value. Red and blue arrows on the color bar indicate significant t-stat value. Female WT: n = 6 mice. Female *Cln3*-/-: n = 7 mice. ***B and C,*** scatter and line plots of N1 amplitude (Ch21) for female WT and *Cln3*-/- mice. Individual data points represent each animal and error bars represent mean ± SEM. Repeated measures two-way ANOVA showed a significant genotype effect (p < 0.0001, with a progressive reduction in N1 in female *Cln3*-/- mice compared to female WT mice. Age-matched genotype comparison showed that female *Cln3*-/- mice exhibited reductions in N1 across age (3 month: p = 0.0016, 5 month: p = 0.0458, 7 month: p = 0.0095, 9 month: p = 0.0001). ***D,*** Trial and subject averaged standard AEP waveforms for female *Cln3*-/- mice from 3 to 9 months of age (Ch21). The vertical dashed line indicates stimulus onset. Shaded areas represent the standard error of the mean (SEM). Blue: 3 month; Cyan: 5 month, Yellow: 7 month, Red: 9 month. The black horizontal lines indicate the time window during which AEP at 9 months is significantly reduced compared to AEP at 3 months (p < 0.05, FDR corrected unpaired t-tests).

Spatiotemporal analysis revealed that between 3 and 7 months, significant differences were primarily localized around the N1 time window (∼40 ms) (p < 0.05, FDR-corrected unpaired t-tests), suggesting early and sustained impairment of thalamocortical auditory processing (Fig. 8A). At 9 months of age, female *Cln3*-/-mice exhibited more pronounced reductions in AEP responses spanning 30-60 ms (N1) and extending to later components (∼300 ms) (p < 0.05, FDR-corrected unpaired t-tests), indicating progressive worsening of auditory processing deficits. This temporal pattern differs from male *Cln3-/-* mice, which show a delayed onset of N1 impairment (Fig. 7).

N1 peak analysis confirmed the results from the spatiotemporal analysis. Female *Cln3-/-* mice showed a consistent reduction in N1 amplitude across all ages (Fig. 8B and 8C). Two-way ANOVA revealed a significant genotype effect (F(1, 44) = 38.37, p < 0.0001, η²_p_ = 0.466) (Fig. 8C). Compared with age-matched WT mice, female *Cln3-/-* mice exhibited significantly reduced N1 amplitude at every time point (3 month: p = 0.0016; 5 month: p = 0.0458; 7 month: p = 0.0095; 9 month: p = 0.0001, Tukey’s test) (Fig. 8B and 8C).

Age also showed a significant effect across pooled genotypes (F(3, 44) = 4.697, p = 0.0062, η²_p_ = 0.243, two-way ANOVA) (Fig. 8B and 8C). While female WT mice showed no significant age-related N1 reduction (p > 0.05, Tukey’s test), female *Cln3-/-* mice exhibited significant N1 reduction at 9 months compared with both 3 and 5 months (3 month vs 9 month: p = 0.0313; 5 month vs 9 month: p = 0.0057, Tukey’s test).

Whole-waveform analysis further confirmed age-dependent attenuation of auditory responses, with significant reductions at 9 months between 30–60 ms and around 200 ms compared with 3 months (p < 0.05, FDR-corrected unpaired t-tests) (Fig. 8D).

Additional analyses of AEP responses to deviant stimuli (Supplementary Fig. 10) and responses at 800 ms ISI (Supplementary Fig. 11) in female mice showed consistent genotype- and age-dependent differences, supporting the robustness of these findings. Together, these results demonstrate that female *Cln3-/-* mice exhibit earlier onset and more pronounced progression of N1 deficits compared with males, indicating sex-dependent vulnerability of auditory thalamocortical function.

### 3.5 Auditory neurophysiological deficits map onto thalamocortical lysosomal storage pathology

To determine how regional lysosomal storage pathology relates to auditory circuit dysfunction, we applied partial least squares (PLS) regression to integrate histological and electrophysiological measures in *Cln3-/-* mice. SCMAS accumulation was quantified across major nodes of the auditory thalamocortical circuit, including the MGv, auditory TRN, and superficial and deep layers of A1. These measures were evaluated in relation to two complementary electrophysiological readouts—N1 amplitude, indexing early thalamocortical processing, and auditory duration MMN, reflecting higher-order auditory function (Ding et al., 2025). Because the datasets were obtained from separate cohorts, analyses were conducted using age- and sex-matched group summaries, enabling identification of latent relationships between distributed pathology and auditory responses.

We first examined how regional lysosomal storage patterns relate to variability in the early AEP component (N1). Both male and female *Cln3*-/- mice displayed age-related reductions in N1 amplitude (Fig. 9A). Among all regions, MGv showed a strong negative coefficient (b_1_ = -3.33, 95% CI [-6.26, -0.03]) with high sign stability (97.63%) (Fig. 9B), indicating that greater thalamic relay pathology robustly predicts reduced N1 amplitude.

**Fig. 9.**
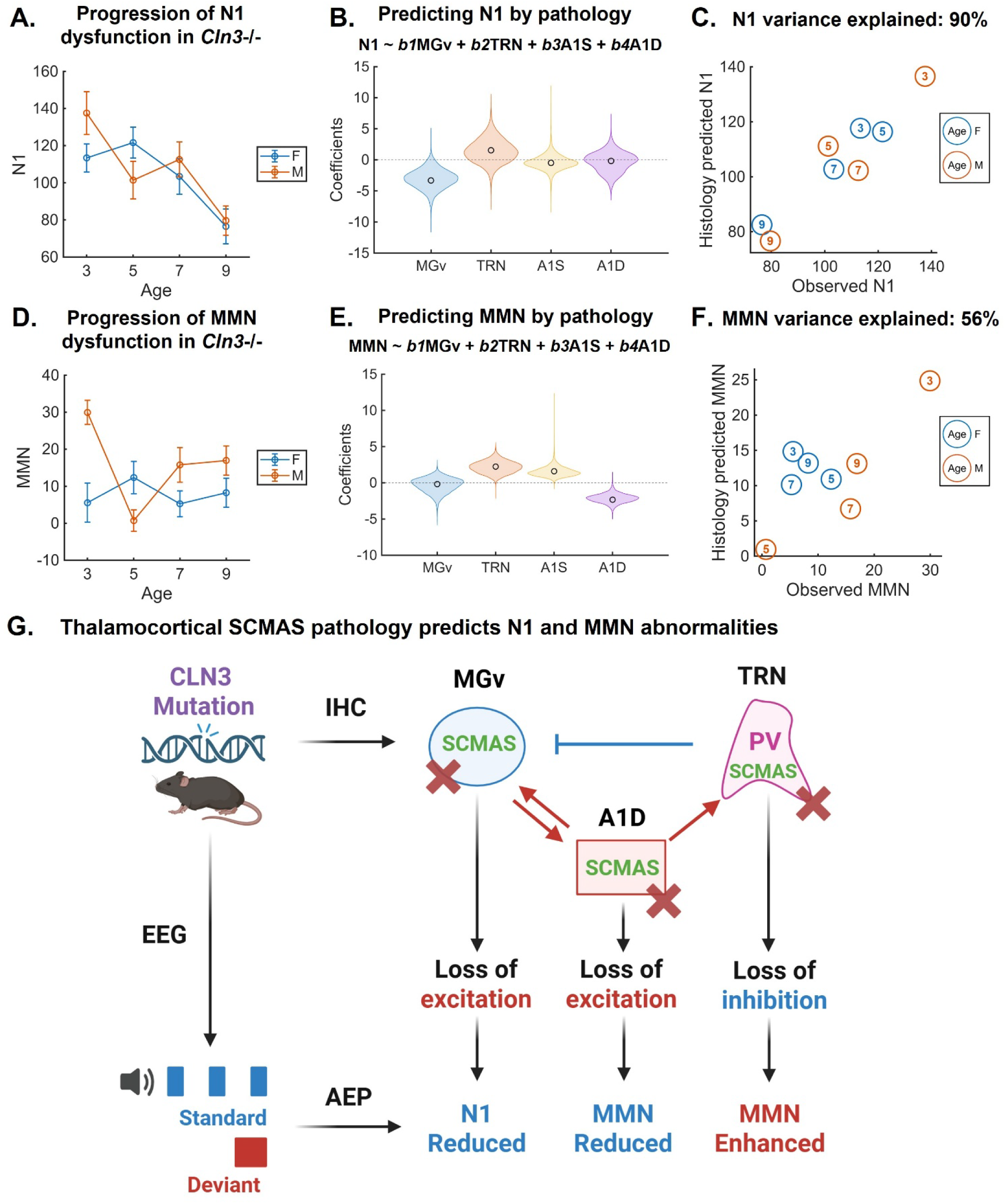
Auditory neurophysiological deficits map onto thalamocortical lysosomal storage pathology in *Cln3-/-* mice. ***A*,** N1 amplitude (Ch21) line plots for male and female *Cln3*-/- mice. Individual data points are group mean ± SEM. N1 progressively reduced across age in both male and female *Cln3*-/- mice. ***B,*** violin plots of the N1 PLS regression coefficients for the TRN, MGv, A1S and A1D. Regression coefficients (b) and corresponding CI were reported in the Results. ***C,*** histology predicted versus observed N1 values in male and female *Cln3*-/- mice. PLS model explained 90% of the N1 variability. ***D*,** auditory duration MMN (Ch21) line plots for male and female *Cln3*-/- mice. Individual data points are group mean ± SEM. *Cln3*-/- mice showed age- and sex-dependent trajectories of auditory duration MMN. ***E,*** violin plots of the MMN PLS regression coefficients for the TRN, MGv, A1S and A1D. Regression coefficients (b) and corresponding CI were reported in the Results. ***F,*** histology predicted versus observed auditory duration MMN values in male and female *Cln3*-/- mice. PLS model explained 56% of the auditory duration MMN variability. ***G,*** summary diagram for the age- and sex-dependent changes in thalamocortical lysosomal storage pathology and auditory neurophysiological responses in the *Cln3*-/- mouse model. Auditory neurophysiological deficits map onto thalamocortical lysosomal storage pathology in *Cln3-/-* mice. MGv SCMAS pathology predicts N1 reduction, TRN SCMAS pathology predicts MMN enhancement, A1D SCMAS pathology predicts MMN reduction.

TRN exhibited a positive coefficient (b_2_ = 1.54, 95% CI [-2.03, 5.35]) with moderate sign stability (78.35%), suggesting a secondary and less consistent contribution (Fig. 9B). In contrast, both A1 superficial (b_3_ = -0.49, 95% CI [-2.68, 1.96], sign stability 68.38%) and deep layers (b_4_ = -0.18, 95% CI [-3.62, 2.56], sign stability 55.26%) showed coefficients near zero with nearly random signs, indicating limited contributions to N1 variability (Fig. 9B).

The observed-versus-predicted relationship demonstrated strong agreement across age × sex groups, with values closely aligned along the unity line and approximately 90% of variance explained (Fig. 9C). These results identify MGv pathology as the dominant anatomical predictor of N1 variability, linking early thalamocortical dysfunction to lysosomal storage burden.

We next examined how lysosomal storage patterns relate to variability in higher-order auditory processing, as indexed by auditory duration MMN. While male *Cln3*-/- mice showed an initial increase, followed by a decline, then a later recovery in the auditory duration MMN, female *Cln3*-/- mice displayed a persistently low level of MMN (Ding et al., 2025) (Fig. 9D).

The PLS model revealed a distinct and distributed anatomical signature associated with MMN (Fig. 9E). In contrast to N1, MGv showed a weak and unstable contribution to MMN (b_1_ = −0.19, 95% CI [−2.55, 1.28], sign stability 58.42%). TRN exhibited a robust positive coefficient (b_2_ = 2.25, 95% CI [0.63, 3.83]) with high sign stability (99.82%), indicating that greater TRN pathology is associated with enhanced MMN responses. A1 superficial layers also showed a positive contribution (b_3_ = 1.60, 95% CI [0.54, 3.05], sign stability 99.95%), whereas A1 deep layers exhibited a strong negative coefficient (b_4_ = -2.33, 95% CI [-3.36, -0.90], sign stability 99.84%), indicating that greater deep-layer A1 pathology predicts reduced MMN.

The PLS model explained approximately 56% of MMN variance across group means (Fig. 9F), indicating a moderate but meaningful association between distributed pathology in thalamocortical circuits and higher-order auditory processing. Together, these findings reveal outcome-specific anatomical signatures linking lysosomal storage pathology to distinct levels of auditory processing: N1 variability is primarily explained by pathology in the thalamic relay (MGv), whereas MMN reflects distributed and opposing contributions across inhibitory (TRN) and cortical (A1) components of the auditory thalamocortical circuit.

## 4. Discussion

This study establishes a circuit-level framework linking lysosomal storage pathology to auditory neurophysiological dysfunction in the *Cln3-/-* mouse model. By integrating histological and electrophysiological analyses, we demonstrate that region- and cell-type–specific SCMAS accumulation maps onto distinct components of auditory processing, revealing how pathology propagates across the auditory thalamocortical circuit.

Our findings identify the MGv as a key locus linking pathology to early auditory deficits. The observation that SCMAS accumulation in the MGv predicts reduced N1 amplitude is consistent with its role as the primary thalamic relay to A1 and a major driver of early central auditory responses (Lee, 2013; Modi and Sahin, 2017). In contrast, higher-order auditory responses (MMN) reflect a distributed and region-specific pattern of pathological influence, involving TRN, MGv, and A1 cortical layers (Naatanen et al., 2007; Lakatos et al., 2020).

These differential anatomical contributions suggest that lysosomal storage pathology disrupts excitation–inhibition (E/I) balance across the auditory thalamocortical circuit. The auditory TRN, enriched in PV+ inhibitory interneurons, provides fast, feedforward inhibition critical for thalamocortical gain control (Li et al., 2020; Takata, 2020). Our findings suggest that pathology in the auditory TRN may weaken inhibitory control over MGv neurons, producing disinhibition that enhances deviant responses. In parallel, pathology in excitatory neurons of the MGv and A1 deep layers (Yuan et al., 2011; Tomioka et al., 2023) may reduce thalamocortical drive and corticothalamic feedback, thereby dampening higher-order auditory responses such as MMN.

These circuit-level effects extend prior studies showing that lysosomal dysfunction disrupts synaptic transmission and neural excitability in CLN3 disease (Grunewald et al., 2017; Ahrens-Nicklas et al., 2019; Gomez-Giro et al., 2019). In other lysosomal storage disorder models, impaired lysosomal function has been linked to mitochondrial dysfunction, oxidative stress, and disturbed Ca²⁺ homeostasis, all of which can compromise neuronal signaling (Ballabio and Gieselmann, 2009; Plotegher and Duchen, 2017; Stepien et al., 2020). Future studies combining circuit-specific manipulations with in-vivo recordings will be critical for establishing causal links between lysosomal pathology and auditory circuit dysfunction.

Our results also reveal evidence for sex-dependent differences in disease progression and circuit compensation. Male Cln3-/- mice exhibited a delayed onset of N1 deficits, emerging between 5 and 9 months, whereas auditory duration MMN showed a more complex trajectory with early enhancement, transient reduction, and partial recovery (Ding et al., 2025). This dissociation suggests that higher-order auditory networks may transiently compensate for early deficits in thalamocortical processing. Such compensation has been observed in other sensory systems, where higher-order cortical areas maintain function despite degradation of primary inputs (Moore and Armstrong, 2003; de Villers-Sidani et al., 2010; Gilbert and Li, 2013; Keck et al., 2013). Because MMN depends on distributed cortical and associative networks, including frontal and parietal regions (Naatanen et al., 2007; Garrido et al., 2009; Pruvost-Robieux et al., 2022), these circuits may support adaptive responses that partially preserve auditory change detection in early disease stages. Future studies targeting higher-order auditory and associative regions will be important to define the mechanisms underlying this compensatory plasticity.

In contrast, female *Cln3-/-* mice exhibited earlier and more sustained impairments in both N1 and MMN, indicating heightened vulnerability of auditory circuits. These findings parallel clinical observations that female CLN3 patients often show more rapid disease progression (Cialone et al., 2012; Nielsen and Ostergaard, 2013). One potential contributor to this sex difference is hormonal modulation of neural and metabolic processes. Sex steroids such as estrogen regulate glutamatergic transmission, GABAergic plasticity, inflammatory signaling, and mitochondrial function (Qiu et al., 2018; Gross and Mermelstein, 2020; Uddin et al., 2020; Mishra et al., 2023). In other neurodegenerative models, including Alzheimer’s disease, estrogen deficiency exacerbates cognitive and behavioral impairments (Liu et al., 2024). Although hormonal status was not assessed in this study, these mechanisms may contribute to the observed sex-specific vulnerability and warrant further investigation.

Importantly, our findings have direct translational implications for biomarker development. Early AEP components, including the N1, reflect the integrity of primary sensory processing and have been widely used as biomarkers in neurodevelopmental and neurodegenerative disorders (Damaschke et al., 2005; Foxe et al., 2011; Paiva et al., 2016; Modi and Sahin, 2017; Sysoeva et al., 2020; Saby et al., 2023). The progressive reduction in N1 amplitude observed in *Cln3-/-*mice suggests a measurable and longitudinally trackable marker of disease progression. Preliminary data from patients with CLN3 disease show a similar trajectory, with initially enhanced AEP responses followed by progressive attenuation with age (Bojanek et al., 2025). The convergence of mouse and human findings supports N1 as a cross-species biomarker of thalamocortical dysfunction. Moreover, reduced N1 amplitude in patients correlates with cognitive impairments, including deficits in verbal intelligence and working memory (Bojanek et al., 2025)., suggesting a potential link between sensory circuit dysfunction and cognitive decline.

AEP and MMN provide complementary windows into auditory processing and together offer a powerful translational framework. AEPs reflect time-locked responses generated by specific neural pathways, allowing relatively direct interpretation of underlying circuitry (Damaschke et al., 2005; Goffin et al., 2014; Modi and Sahin, 2017; Pruvost-Robieux et al., 2022). In contrast, MMN reflects higher-order auditory change detection (Molholm et al., 2005; Garrido et al., 2009; Ross and Hamm, 2020). By subtracting the standard waveform from the deviant waveform, MMN reduces background variation and provides a robust measure of neural processes distributed across cortical networks (Naatanen et al., 2007; Fitzgerald and Todd, 2020). The combined use of these measures enables simultaneous assessment of primary and higher-order auditory function. Deficits in AEP and MMN have been reported in disorders such as Alzheimer’s disease and schizophrenia (Foxe et al., 2011; Nagai et al., 2013; Francisco et al., 2020; Du et al., 2025; Bayat et al., 2026), highlighting their sensitivity to circuit dysfunction. In CLN3 disease, the integration of these measures captures both early thalamocortical impairment and higher-order network dysfunction, providing a comprehensive framework for disease monitoring and therapeutic evaluation.

Several limitations should be considered. First, EEG signals reflect large-scale population activity and do not provide precise anatomical localization of signal sources (Sur and Sinha, 2009; Light et al., 2010). Combining EEG with circuit-specific recording or manipulation approaches will be necessary to resolve regional contributions more precisely. Second, MMN reflects distributed processing involving frontal and parietal regions beyond the auditory thalamocortical circuit (Molholm et al., 2005; Lakatos et al., 2020). Future studies incorporating these associative regions will be important for fully characterizing higher-order dysfunction. Finally, although we identify strong associations between SCMAS accumulation and neurophysiological changes, these relationships remain correlative, and causal mechanisms require direct experimental testing.

## 5. Conclusion

In summary, this study demonstrates that lysosomal storage pathology in *Cln3-/-* mice exhibits region-, cell type–, and sex-specific patterns that map onto distinct auditory neurophysiological deficits. By linking thalamocortical SCMAS accumulation to functional deficits, this work establishes a circuit-level framework for understanding central auditory dysfunction in CLN3 disease. These findings provide a foundation for mechanistic investigations of circuit vulnerability and support the development of translational EEG-based biomarkers for disease progression and therapeutic intervention.

## Declarations

### Ethics approval and consent to participate

Experiments were conducted in accordance with ethical standards for the care and use of animals of the University Committee on Animal Resource (UCAR) at the University of Rochester Medical Center (URMC, NY)

### Consent for publication

All authors have provided consent for publication

### Data Availability

The data used in the current study will be available upon reasonable request

### Authors’ contributions

YD: Writing - Original Draft, Conceptualization, Methodology, Software, Formal analysis, Investigation, Visualization, Data Curation. JF: Writing - Review & Editing, Conceptualization, Methodology, Software, Resources, Investigation. VP: Writing - Review & Editing, Methodology, Resources. GR: Writing - Review & Editing, Resources, Investigation. AGS: Writing - Review & Editing, Resources. HEC: Writing - Review & Editing, Resources. SAS: Writing - Review & Editing, Resources. EGF: Writing - Review & Editing, Conceptualization, Validation, Project administration, Supervision. JJF: Writing - Review & Editing, Conceptualization, Validation, Supervision, Project administration, Funding. KHW: Writing - Review & Editing, Project administration, Conceptualization, Methodology, Validation, Supervision, Funding.

## Acknowledgments

We thank members of Wang Lab and the Frederick J. and Marion A. Schindler Cognitive Neurophysiology Lab for critical discussions. We acknowledge the use of the Center for Musculoskeletal Research (CMSR) and the Center for Advanced Microscopy & Nanoscopy (CALMN) at University of Rochester Medical Center for imaging service and support. These facilities are supported by the National Institutes of Health (AR069655, P50 HD103536).

## Reference

Adams HR, Mink JW, University of Rochester Batten Center Study G (2013) Neurobehavioral features and natural history of juvenile neuronal ceroid lipofuscinosis (Batten disease). J Child Neurol 28:1128–1136.

Adams HR, Kwon J, Marshall FJ, de Blieck EA, Pearce DA, Mink JW (2007) Neuropsychological symptoms of juvenile-onset batten disease: experiences from 2 studies. J Child Neurol 22:621–627.

Ahrens-Nicklas RC, Tecedor L, Hall AF, Lysenko E, Cohen AS, Davidson BL, Marsh ED (2019) Neuronal network dysfunction precedes storage and neurodegeneration in a lysosomal storage disorder. JCI Insight 4.

Appler JM, Goodrich LV (2011) Connecting the ear to the brain: Molecular mechanisms of auditory circuit assembly. Prog Neurobiol 93:488–508.

Ballabio A, Gieselmann V (2009) Lysosomal disorders: from storage to cellular damage. Biochim Biophys Acta 1793:684–696.

Banach-Petrosky W, Larrimore KE, Sleat EH, Bazer A, Samuels B, Tan Y, Melton AC, Ichida JK, Logan TP, Lobel P, Sleat DE (2025) Elevated tripeptidyl-peptidase 1 corrects multiple disease phenotypes in a mouse model of juvenile neuronal ceroid lipofuscinosis. Mol Ther Methods Clin Dev 33:101587.

Bayat A, Mirmomeni G, Aiken S, Jafari Z (2026) Meta-Analyses of Auditory Evoked Potentials as Alzheimer Biomarkers. Ear Hear 47:95–106.

Beebe NL, Mellott JG, Schofield BR (2018) Inhibitory Projections from the Inferior Colliculus to the Medial Geniculate body Originate from Four Subtypes of GABAergic Cells. eNeuro 5.

Bellettato CM, Scarpa M (2010) Pathophysiology of neuropathic lysosomal storage disorders. J Inherit Metab Dis 33:347–362.

Bennett MJ, Hofmann SL (1999) The neuronal ceroid-lipofuscinoses (Batten disease): a new class of lysosomal storage diseases. J Inherit Metab Dis 22:535–544.

Biacabe B, Chevallier JM, Avan P, Bonfils P (2001) Functional anatomy of auditory brainstem nuclei: application to the anatomical basis of brainstem auditory evoked potentials. Auris Nasus Larynx 28:85–94.

Bojanek EK, Lang ER, Adams HR, Vermilion J, Augustine EF, Brima T, Nasimjonova S, Freedman EG, Foxe JJ (2025) Longitudinal Exploration of Auditory Sensory-Perceptual Processing in CLN3 Disease (Juvenile Neuronal Ceroid Lipofuscinosis (Batten disease)): A High-Density Auditory Evoked Potential (AEP) Study. bioRxiv:2025.2011.2019.689311.

Brima T, Freedman EG, Prinsloo KD, Augustine EF, Adams HR, Wang KH, Mink JW, Shaw LH, Mantel EP, Foxe JJ (2024) Assessing the integrity of auditory sensory memory processing in CLN3 disease (Juvenile Neuronal Ceroid Lipofuscinosis (Batten disease)): an auditory evoked potential study of the duration-evoked mismatch negativity (MMN). J Neurodev Disord 16:3.

Burkovetskaya M, Karpuk N, Kielian T (2017) Age-dependent alterations in neuronal activity in the hippocampus and visual cortex in a mouse model of Juvenile Neuronal Ceroid Lipofuscinosis (CLN3). Neurobiol Dis 100:19–29.

Centa JL, Stratton MP, Pratt MA, Osterlund Oltmanns JR, Wallace DG, Miller SA, Weimer JM, Hastings ML (2023) Protracted CLN3 Batten disease in mice that genetically model an exon-skipping therapeutic approach. Mol Ther Nucleic Acids 33:15–27.

Cialone J, Adams H, Augustine EF, Marshall FJ, Kwon JM, Newhouse N, Vierhile A, Levy E, Dure LS, Rose KR, Ramirez-Montealegre D, de Blieck EA, Mink JW (2012) Females experience a more severe disease course in Batten disease. J Inherit Metab Dis 35:549–555.

Clemente-Perez A, Makinson SR, Higashikubo B, Brovarney S, Cho FS, Urry A, Holden SS, Wimer M, David C, Fenno LE, Acsady L, Deisseroth K, Paz JT (2017) Distinct Thalamic Reticular Cell Types Differentially Modulate Normal and Pathological Cortical Rhythms. Cell Rep 19:2130–2142.

Cohen J (1973) Eta-Squared and Partial Eta-Squared in Fixed Factor Anova Designs. Educ Psychol Meas 33:107–112.

Cohen J (1988) Statistical Power Analysis for the Behavioral-Sciences - Cohen,J. Percept Motor Skill 67:1007–1007.

Consortium TIBD (1995) Isolation of a novel gene underlying Batten disease, CLN3. The International Batten Disease Consortium. Cell 82:949–957.

Damaschke J, Riedel H, Kollmeier B (2005) Neural correlates of the precedence effect in auditory evoked potentials. Hear Res 205:157–171.

de Villers-Sidani E, Alzghoul L, Zhou X, Simpson KL, Lin RC, Merzenich MM (2010) Recovery of functional and structural age-related changes in the rat primary auditory cortex with operant training. Proc Natl Acad Sci U S A 107:13900–13905.

Ding SL, Tecedor L, Stein CS, Davidson BL (2011) A knock-in reporter mouse model for Batten disease reveals predominant expression of Cln3 in visual, limbic and subcortical motor structures. Neurobiol Dis 41:237–248.

Ding Y, Feng J, Prifti V, Rico GA, Solorzano AG, Chang HE, Freedman EG, Foxe JJ, Wang KH (2025) Sex-specific and age-related progression of auditory neurophysiological deficits in the Cln3 mouse model of Batten disease. J Neurodev Disord 17:67.

Doucet JR, Rose L, Ryugo DK (2002) The cellular origin of corticofugal projections to the superior olivary complex in the rat. Brain Res 925:28–41.

Du L, Jiao X, Shi K, Mo D, Zhou J, Shi Y, Zhang X, Zeng Z, Li H, Duan D, Tong S, Sun J, Cui D (2025) Mismatch negativity deficits in schizophrenia and bipolar disorder. J Psychiatr Res 188:94–103.

Elleder M, Sokolova J, Hrebicek M (1997) Follow-up study of subunit c of mitochondrial ATP synthase (SCMAS) in Batten disease and in unrelated lysosomal disorders. Acta Neuropathol 93:379–390.

Elmerskog B, Tossebro AG, Atkinson R, Rokne S, Cole B, Ockelford A, Adams HR (2020) Overview of advances in educational and social supports for young persons with NCL disorders. Biochim Biophys Acta Mol Basis Dis 1866:165480.

Fitzgerald K, Todd J (2020) Making Sense of Mismatch Negativity. Front Psychiatry 11:468.

Foxe JJ, Yeap S, Snyder AC, Kelly SP, Thakore JH, Molholm S (2011) The N1 auditory evoked potential component as an endophenotype for schizophrenia: high-density electrical mapping in clinically unaffected first-degree relatives, first-episode, and chronic schizophrenia patients. Eur Arch Psychiatry Clin Neurosci 261:331–339.

Francisco AA, Foxe JJ, Horsthuis DJ, DeMaio D, Molholm S (2020) Assessing auditory processing endophenotypes associated with Schizophrenia in individuals with 22q11.2 deletion syndrome. Transl Psychiatry 10:85.

Garrido MI, Kilner JM, Stephan KE, Friston KJ (2009) The mismatch negativity: a review of underlying mechanisms. Clin Neurophysiol 120:453–463.

Gilbert CD, Li W (2013) Top-down influences on visual processing. Nat Rev Neurosci 14:350–363.

Goffin D, Brodkin ES, Blendy JA, Siegel SJ, Zhou Z (2014) Cellular origins of auditory event-related potential deficits in Rett syndrome. Nat Neurosci 17:804–806.

Gomez-Giro G, Arias-Fuenzalida J, Jarazo J, Zeuschner D, Ali M, Possemis N, Bolognin S, Halder R, Jager C, Kuper WFE, van Hasselt PM, Zaehres H, Del Sol A, van der Putten H, Scholer HR, Schwamborn JC (2019) Synapse alterations precede neuronal damage and storage pathology in a human cerebral organoid model of CLN3-juvenile neuronal ceroid lipofuscinosis. Acta Neuropathol Commun 7:222.

Gonsalvez CL, Polich J (2002) P300 amplitude is determined by target-to-target interval. Psychophysiology 39:388–396.

Gross KS, Mermelstein PG (2020) Estrogen receptor signaling through metabotropic glutamate receptors. Vitam Horm 114:211–232.

Grunewald B, Lange MD, Werner C, O’Leary A, Weishaupt A, Popp S, Pearce DA, Wiendl H, Reif A, Pape HC, Toyka KV, Sommer C, Geis C (2017) Defective synaptic transmission causes disease signs in a mouse model of juvenile neuronal ceroid lipofuscinosis. Elife 6.

Hall NA, Lake BD, Dewji NN, Patrick AD (1991) Lysosomal storage of subunit c of mitochondrial ATP synthase in Batten’s disease (ceroid-lipofuscinosis). Biochem J 275 ( Pt 1):269–272.

Homma NY, Bajo VM (2021) Lemniscal Corticothalamic Feedback in Auditory Scene Analysis. Front Neurosci 15:723893.

Hu H, Gan J, Jonas P (2014) Interneurons. Fast-spiking, parvalbumin(+) GABAergic interneurons: from cellular design to microcircuit function. Science 345:1255263.

Jana M, Dutta D, Poddar J, Pahan K (2023) Activation of PPARalpha Exhibits Therapeutic Efficacy in a Mouse Model of Juvenile Neuronal Ceroid Lipofuscinosis. J Neurosci 43:1814–1829.

Johnson TB, Cain JT, White KA, Ramirez-Montealegre D, Pearce DA, Weimer JM (2019) Therapeutic landscape for Batten disease: current treatments and future prospects. Nat Rev Neurol 15:161–178.

Keck T, Keller GB, Jacobsen RI, Eysel UT, Bonhoeffer T, Hubener M (2013) Synaptic scaling and homeostatic plasticity in the mouse visual cortex in vivo. Neuron 80:327–334.

Kim WD, Wilson-Smillie M, Thanabalasingam A, Lefrancois S, Cotman SL, Huber RJ (2022) Autophagy in the Neuronal Ceroid Lipofuscinoses (Batten Disease). Front Cell Dev Biol 10:812728.

Lakatos P, O’Connell MN, Barczak A, McGinnis T, Neymotin S, Schroeder CE, Smiley JF, Javitt DC (2020) The Thalamocortical Circuit of Auditory Mismatch Negativity. Biol Psychiatry 87:770–780.

Lakens D (2013) Calculating and reporting effect sizes to facilitate cumulative science: a practical primer for -tests and ANOVAs. Front Psychol 4.

Lee CC (2013) Thalamic and cortical pathways supporting auditory processing. Brain Lang 126:22–28.

Li Y et al. (2020) Distinct subnetworks of the thalamic reticular nucleus. Nature 583:819–824.

Light GA, Williams LE, Minow F, Sprock J, Rissling A, Sharp R, Swerdlow NR, Braff DL (2010) Electroencephalography (EEG) and event-related potentials (ERPs) with human participants. Curr Protoc Neurosci Chapter 6:Unit 6 25 21–24.

Liu F, Liu Y, Shen X, Du J, Zhang H, Hou X (2024) Ovariectomy exacerbates the disturbance of excitation-inhibition balance in the brain of APP/PS-1/tau mice. Front Mol Neurosci 17:1391082.

Luo Y, Wang L, Cao Y, Shen Y, Gu Y, Wang L (2024) Reduced excitatory activity in the developing mPFC mediates a PV(H)-to-PV(L) transition and impaired social cognition in autism spectrum disorders. Transl Psychiatry 14:325.

Mazer P, Carneiro F, Domingo J, Pasion R, Silveira C, Ferreira-Santos F (2024) Systematic review and meta-analysis of the visual mismatch negativity in schizophrenia. Eur J Neurosci 59:2863–2874.

Meltzer NE, Ryugo DK (2006) Projections from auditory cortex to cochlear nucleus: A comparative analysis of rat and mouse. Anat Rec A Discov Mol Cell Evol Biol 288:397–408.

Mink JW, Augustine EF, Adams HR, Marshall FJ, Kwon JM (2013) Classification and natural history of the neuronal ceroid lipofuscinoses. J Child Neurol 28:1101–1105.

Mishra P, Davies DA, Albensi BC (2023) The Interaction Between NF-kappaB and Estrogen in Alzheimer’s Disease. Mol Neurobiol 60:1515–1526.

Mitchison HM, Bernard DJ, Greene ND, Cooper JD, Junaid MA, Pullarkat RK, de Vos N, Breuning MH, Owens JW, Mobley WC, Gardiner RM, Lake BD, Taschner PE, Nussbaum RL (1999) Targeted disruption of the Cln3 gene provides a mouse model for Batten disease. The Batten Mouse Model Consortium [corrected]. Neurobiol Dis 6:321–334.

Modi ME, Sahin M (2017) Translational use of event-related potentials to assess circuit integrity in ASD. Nat Rev Neurol 13:160–170.

Mole SE, Mitchison HM, Munroe PB (1999) Molecular basis of the neuronal ceroid lipofuscinoses: mutations in CLN1, CLN2, CLN3, and CLN5. Hum Mutat 14:199–215.

Molholm S, Martinez A, Ritter W, Javitt DC, Foxe JJ (2005) The neural circuitry of pre-attentive auditory change-detection: an fMRI study of pitch and duration mismatch negativity generators. Cereb Cortex 15:545–551.

Moore T, Armstrong KM (2003) Selective gating of visual signals by microstimulation of frontal cortex. Nature 421:370–373.

Morison LD, Whiteman IT, Vogel AP, Tilbrook L, Fahey MC, Braden R, Bredebusch J, Hildebrand MS, Scheffer IE, Morgan AT (2025) Speech, Language and Non-verbal Communication in CLN2 and CLN3 Batten Disease. J Inherit Metab Dis 48:e12838.

Munroe PB, Mitchison HM, O’Rawe AM, Anderson JW, Boustany RM, Lerner TJ, Taschner PE, de Vos N, Breuning MH, Gardiner RM, Mole SE (1997) Spectrum of mutations in the Batten disease gene, CLN3. Am J Hum Genet 61:310–316.

Naatanen R, Paavilainen P, Rinne T, Alho K (2007) The mismatch negativity (MMN) in basic research of central auditory processing: a review. Clin Neurophysiol 118:2544–2590.

Nagai T, Tada M, Kirihara K, Araki T, Jinde S, Kasai K (2013) Mismatch negativity as a "translatable" brain marker toward early intervention for psychosis: a review. Front Psychiatry 4:115.

Nielsen AK, Ostergaard JR (2013) Do females with juvenile ceroid lipofuscinosis (Batten disease) have a more severe disease course? The Danish experience. Eur J Paediatr Neurol 17:265–268.

Ostergaard JR (2016) Juvenile neuronal ceroid lipofuscinosis (Batten disease): current insights. Degener Neurol Neuromuscul Dis 6:73–83.

Paiva TO, Almeida PR, Ferreira-Santos F, Vieira JB, Silveira C, Chaves PL, Barbosa F, Marques-Teixeira J (2016) Similar sound intensity dependence of the N1 and P2 components of the auditory ERP: Averaged and single trial evidence. Clin Neurophysiol 127:499–508.

Pinault D (2004) The thalamic reticular nucleus: structure, function and concept. Brain Res Brain Res Rev 46:1–31.

Platt FM, d’Azzo A, Davidson BL, Neufeld EF, Tifft CJ (2018) Lysosomal storage diseases. Nat Rev Dis Primers 4:27.

Plotegher N, Duchen MR (2017) Mitochondrial Dysfunction and Neurodegeneration in Lysosomal Storage Disorders. Trends Mol Med 23:116–134.

Pontikis CC, Cotman SL, MacDonald ME, Cooper JD (2005) Thalamocortical neuron loss and localized astrocytosis in the Cln3Deltaex7/8 knock-in mouse model of Batten disease. Neurobiol Dis 20:823–836.

Pruvost-Robieux E, Marchi A, Martinelli I, Bouchereau E, Gavaret M (2022) Evoked and Event-Related Potentials as Biomarkers of Consciousness State and Recovery. J Clin Neurophysiol 39:22–31.

Qiu J, Rivera HM, Bosch MA, Padilla SL, Stincic TL, Palmiter RD, Kelly MJ, Ronnekleiv OK (2018) Estrogenic-dependent glutamatergic neurotransmission from kisspeptin neurons governs feeding circuits in females. Elife 7.

Ritter W, Sussman E, Molholm S, Foxe JJ (2002) Memory reactivation or reinstatement and the mismatch negativity. Psychophysiology 39:158–165.

Ross JM, Hamm JP (2020) Cortical Microcircuit Mechanisms of Mismatch Negativity and Its Underlying Subcomponents. Front Neural Circuits 14:13.

Ruden JB, Dugan LL, Konradi C (2021) Parvalbumin interneuron vulnerability and brain disorders. Neuropsychopharmacology 46:279–287.

Saby JN, Peters SU, Benke TA, Standridge SM, Swanson LC, Lieberman DN, Olson HE, Key AP, Percy AK, Neul JL, Nelson CA, Roberts TPL, Marsh ED (2023) Comparison of evoked potentials across four related developmental encephalopathies. J Neurodev Disord 15:10.

Slater BJ, Isaacson JS (2020) Interhemispheric Callosal Projections Sharpen Frequency Tuning and Enforce Response Fidelity in Primary Auditory Cortex. eNeuro 7.

Smith PH, Uhlrich DJ, Manning KA, Banks MI (2012) Thalamocortical projections to rat auditory cortex from the ventral and dorsal divisions of the medial geniculate nucleus. J Comp Neurol 520:34–51.

Stepien KM, Roncaroli F, Turton N, Hendriksz CJ, Roberts M, Heaton RA, Hargreaves I (2020) Mechanisms of Mitochondrial Dysfunction in Lysosomal Storage Disorders: A Review. J Clin Med 9.

Steullet P, Cabungcal JH, Bukhari SA, Ardelt MI, Pantazopoulos H, Hamati F, Salt TE, Cuenod M, Do KQ, Berretta S (2018) The thalamic reticular nucleus in schizophrenia and bipolar disorder: role of parvalbumin-expressing neuron networks and oxidative stress. Mol Psychiatry 23:2057–2065.

Sur S, Sinha VK (2009) Event-related potential: An overview. Ind Psychiatry J 18:70–73.

Suthakar K, Liberman MC (2021) Auditory-nerve responses in mice with noise-induced cochlear synaptopathy. J Neurophysiol 126:2027–2038.

Sysoeva OV, Molholm S, Djukic A, Frey HP, Foxe JJ (2020) Atypical processing of tones and phonemes in Rett Syndrome as biomarkers of disease progression. Transl Psychiatry 10:188.

Takata N (2020) Thalamic reticular nucleus in the thalamocortical loop. Neurosci Res 156:32–40.

Tang C et al. (2021) A human model of Batten disease shows role of CLN3 in phagocytosis at the photoreceptor-RPE interface. Commun Biol 4:161.

Tomioka R, Takemoto M, Song WJ (2023) Neurochemical properties for defining subdivisions of the mouse medial geniculate body. Hear Res 431:108724.

Uddin MS, Rahman MM, Jakaria M, Rahman MS, Hossain MS, Islam A, Ahmed M, Mathew B, Omar UM, Barreto GE, Ashraf GM (2020) Estrogen Signaling in Alzheimer’s Disease: Molecular Insights and Therapeutic Targets for Alzheimer’s Dementia. Mol Neurobiol 57:2654–2670.

Vallesi A, Lozano VN, Correa A (2013) Dissociating temporal preparation processes as a function of the inter-trial interval duration. Cognition 127:22–30.

Wang Y, Ye M, Kuang X, Li Y, Hu S (2018) A simplified morphological classification scheme for pyramidal cells in six layers of primary somatosensory cortex of juvenile rats. IBRO Rep 5:74–90.

Wang Y, Li K, Chen W, Chen C, Yong AJH, Zhang X, Tynan-La Fontaine M, Jan YN, Jan LY (2026) CLN3 mediates chloride efflux from lysosomes. Neuron 114:868–883 e867.

Weimer JM, Custer AW, Benedict JW, Alexander NA, Kingsley E, Federoff HJ, Cooper JD, Pearce DA (2006) Visual deficits in a mouse model of Batten disease are the result of optic nerve degeneration and loss of dorsal lateral geniculate thalamic neurons. Neurobiol Dis 22:284–293.

Yang C, Wang X (2021) Lysosome biogenesis: Regulation and functions. J Cell Biol 220.

Yuan K, Fink KL, Winer JA, Schreiner CE (2011) Local connection patterns of parvalbumin-positive inhibitory interneurons in rat primary auditory cortex. Hear Res 274:121–128.

Zinnamon FA, Harrison FG, Wenas SS, Liu Q, Wang KH, Linden JF (2023) Increased Central Auditory Gain and Decreased Parvalbumin-Positive Cortical Interneuron Density in the Df1/+ Mouse Model of Schizophrenia Correlate With Hearing Impairment. Biol Psychiatry Glob Open Sci 3:386–397.

Znamenskiy P, Zador AM (2013) Corticostriatal neurons in auditory cortex drive decisions during auditory discrimination. Nature 497:482–485.

