## Supplementary figures for "Distinct Auditory Thalamocortical Pathologies Underlie Emerging Neurophysiological Dysfunction in a Cln3 Mouse Model of Batten Disease"

### MGv SCMAS

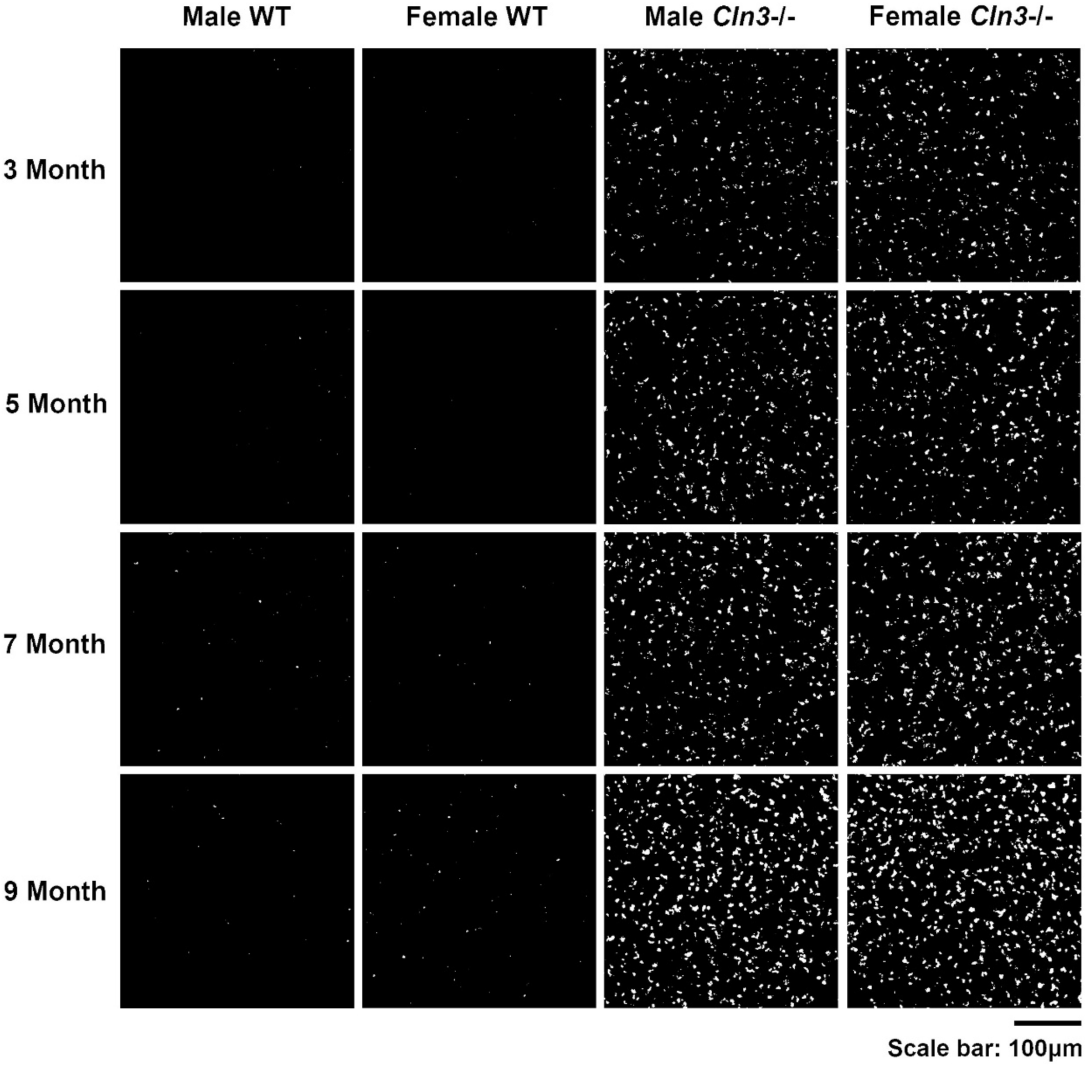
Male WT
Female WT
Male *Cln3*-/-
Female *Cln3*-/-
3 Month
5 Month
7 Month
9 Month
Scale bar: 100μm

**Supplementary Fig. 1 Early sex difference in MGv lysosomal storage accumulation.** Binarized representative images of SCMAS accumulation in the MGv after background subtraction in male WT, female WT, male *Cln3*-/- and female *Cln3*-/- mice between 3 to 9 months of age. Scale bar: 100μm. WT mice of both sexes showed minimal SCMAS accumulation. *Cln3*-/- mice of both sexes exhibited progressive SCMAS accumulation. Female *Cln3*-/- mice showed higher SCMAS accumulation in the MGv at 3 months of age.

### Auditory TRN SCMAS

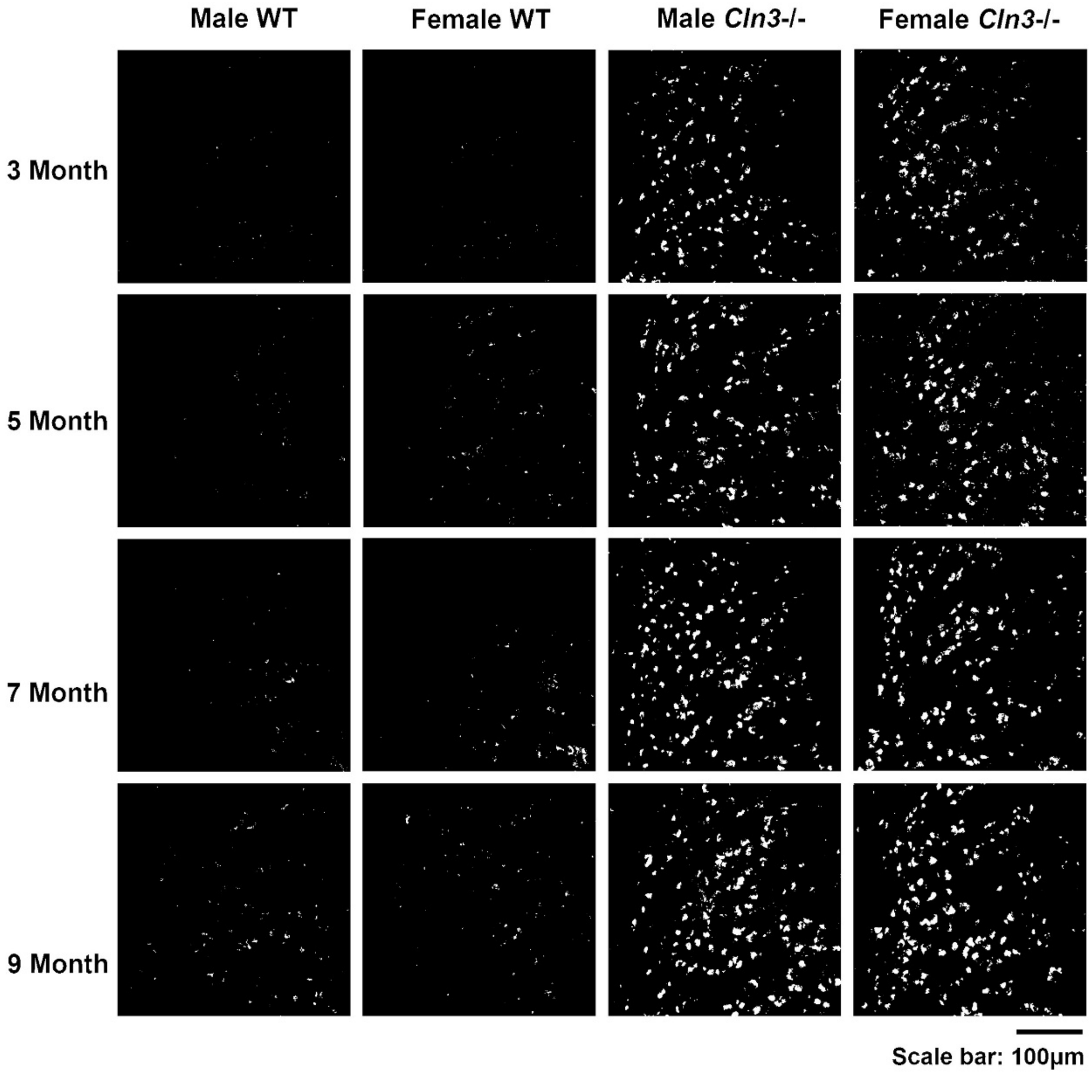

**Supplementary Fig. 2 Late-emerging sex difference in lysosomal storage in the auditory TRN.** Binarized representative images of SCMAS accumulation in the auditory TRN after background subtraction in male WT, female WT, male *Cln3*-/- and female *Cln3*-/- mice between 3 and 9 months of age. Scale bar: 100µm. WT mice of both sexes showed minimal SCMAS accumulation. *Cln3*-/- mice of both sexes exhibited progressive SCMAS accumulation. Female *Cln3*-/- mice showed higher SCMAS accumulation in the auditory TRN at 9 months of age.

### A1 (L1-L4) SCMAS

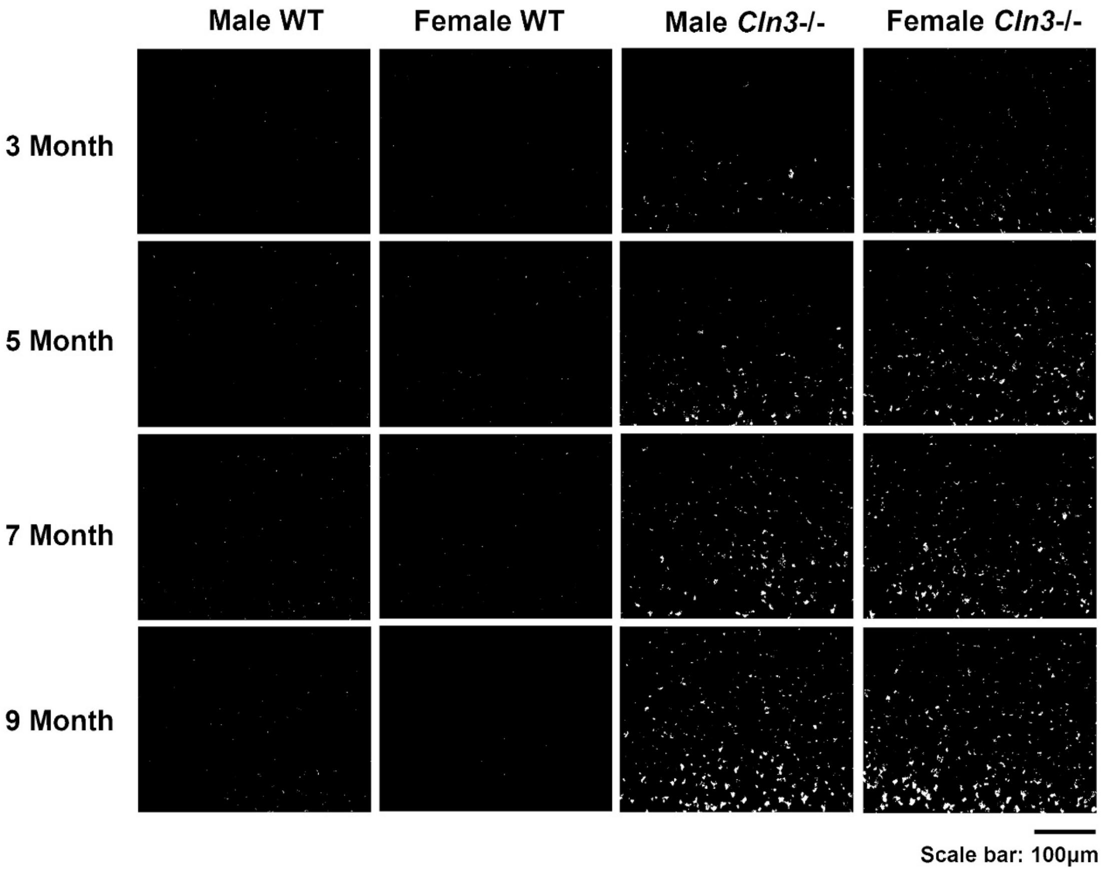

**Supplementary Fig. 3 Lysosomal storage accumulation in A1 superficial layers is primarily driven by genotype and age rather than sex.** Binarized representative images of SCMAS accumulation in the A1 superficial layer after background subtraction in male WT, female WT, male *Cln3*-/- and female *Cln3*-/- mice between 3 and 9 months of age. Scale bar: 100μm. WT mice of both sexes showed minimal SCMAS accumulation. *Cln3*-/- mice of both sexes exhibited progressive SCMAS accumulation. Male and female *Cln3*-/- mice showed no sex-specific difference in SCMAS accumulation in the A1 superficial layers.

### A1 (L5-L6) SCMAS

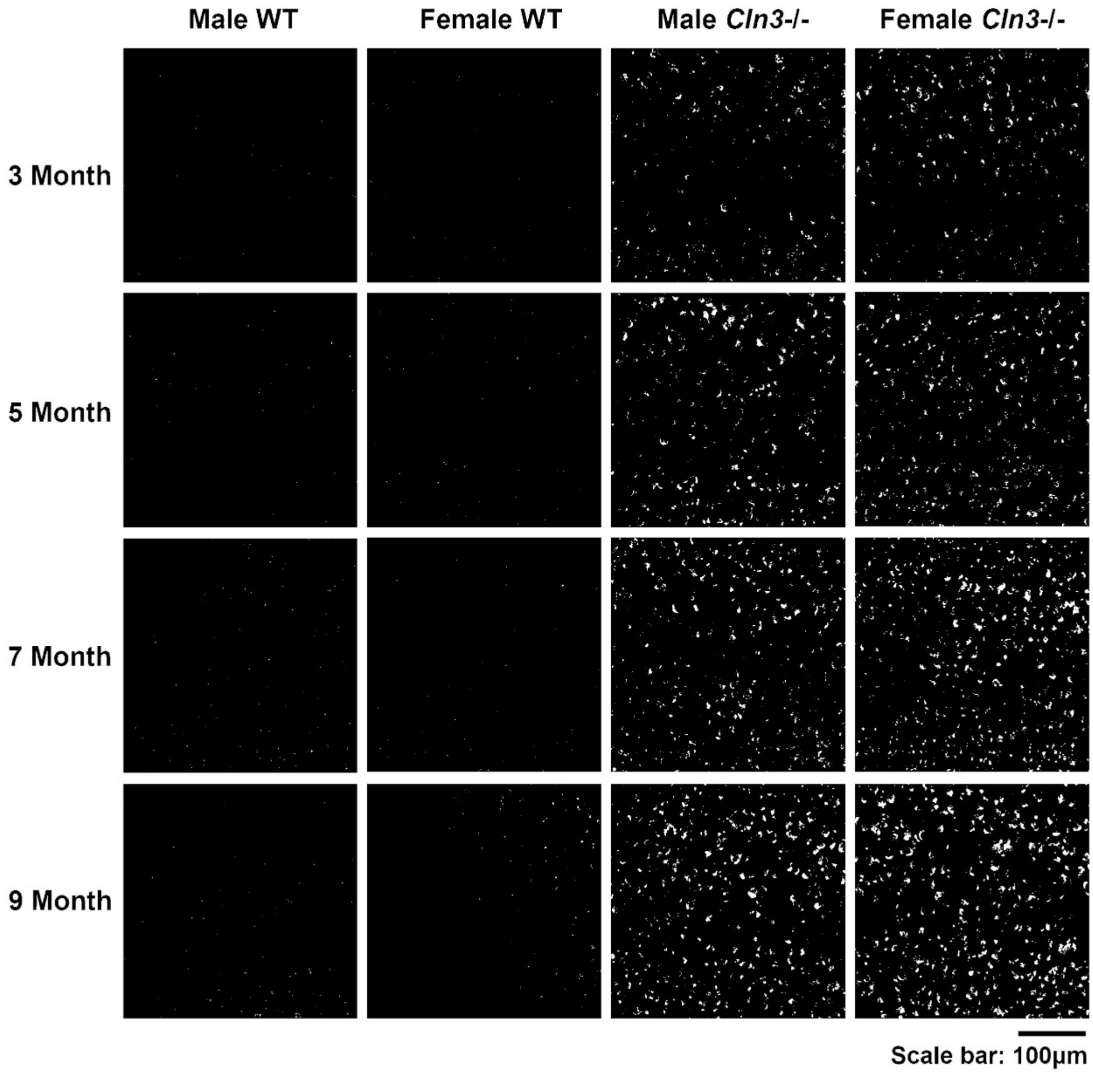

**Supplementary Fig. 4 *Cln3*-/- mice showed age- and sex-dependent SCMAS accumulation in the A1 deep layers.** Binarized representative images of SCMAS accumulation in the A1 deep layer after background subtraction in male WT, female WT, male *Cln3*-/- and female *Cln3*-/- mice between 3 and 9 months of age. Scale bar: 100μm. WT mice of both sexes showed minimal SCMAS accumulation. *Cln3*-/- mice of both sexes exhibited progressive SCMAS accumulation. Female *Cln3*-/- mice showed higher SCMAS accumulation in the A1 deep layers at 3 and 9 months of age. Male *Cln3*-/- mice showed higher SCMAS accumulation in the A1 deep layers at 5 months of age.

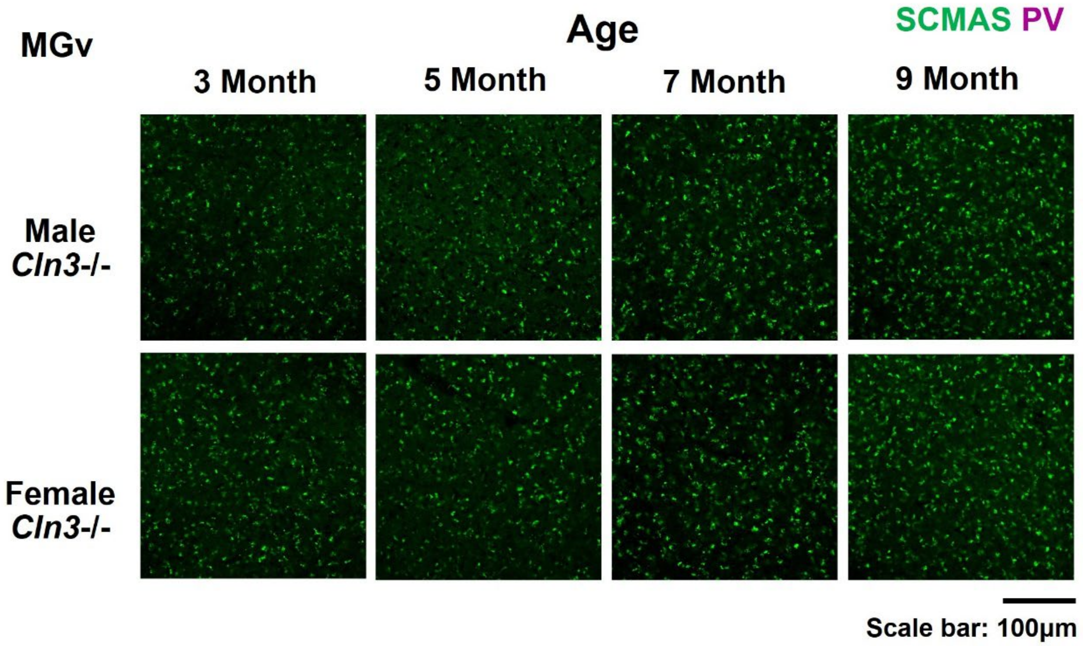

MGv
Age
SCMAS PV
3 Month
5 Month
7 Month
9 Month
Male
Cln3-/-
Female
Cln3-/-
Scale bar: 100µm

**Supplementary Fig. 5 There was no PV and SCMAS co-localization in the MGv in *Cln3*-/- mice.** Magenta: PV; Green: SCMAS. Scale bar: 100μm. Male and female *Cln3*-/- mice showed progressive SCMAS accumulation with no PV+ cells in the MGv between 3 and 9 months of age.

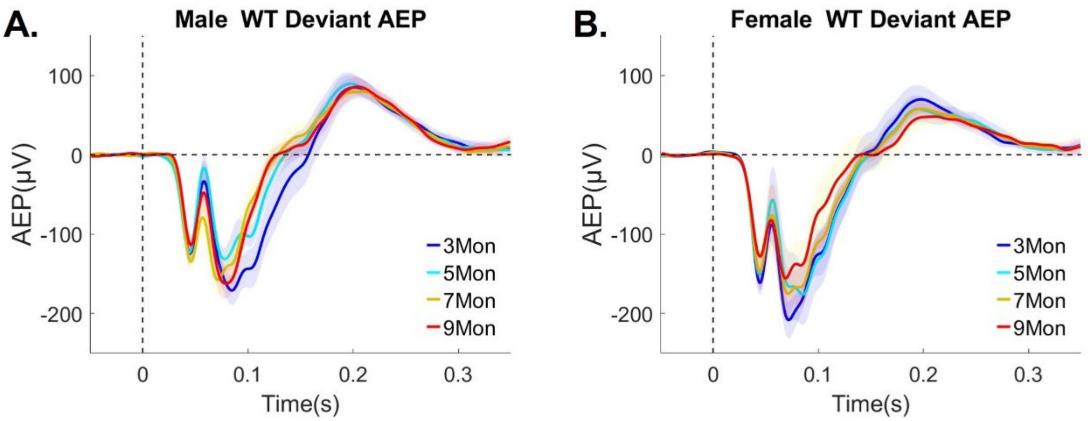

A.
Male WT Deviant AEP
B.
Female WT Deviant AEP
AEP(μV)
100
0
-100
-200
0
0.1
0.2
0.3
Time(s)
3Mon
5Mon
7Mon
9Mon

**Supplementary Fig. 6 WT mice of both sexes showed consistent AEP responses to deviant tones across age.** ***A and B*****,** Trial and subject averaged standard AEP waveforms for male (A) and female (B) WT mice aged 3 to 9 months, recorded from a centrally located channel (Ch21). The ISI is 1600 ms. The vertical dashed line indicates stimulus onset. AEPs in response to standard are presented, with the shaded areas representing the standard error of the mean (SEM). Blue: 3 month; Cyan: 5 month, Yellow: 7 month, Red: 9 month. Male WT: n = 11 mice for 3-, 7- and 9- month-old groups; n = 14 for the 5-month-old group. Female WT: n = 6 mice for all ages. All animals were recorded longitudinally from 3 to 9 months of age, except three additional male mice recorded only at 5 months of age in pilot studies.

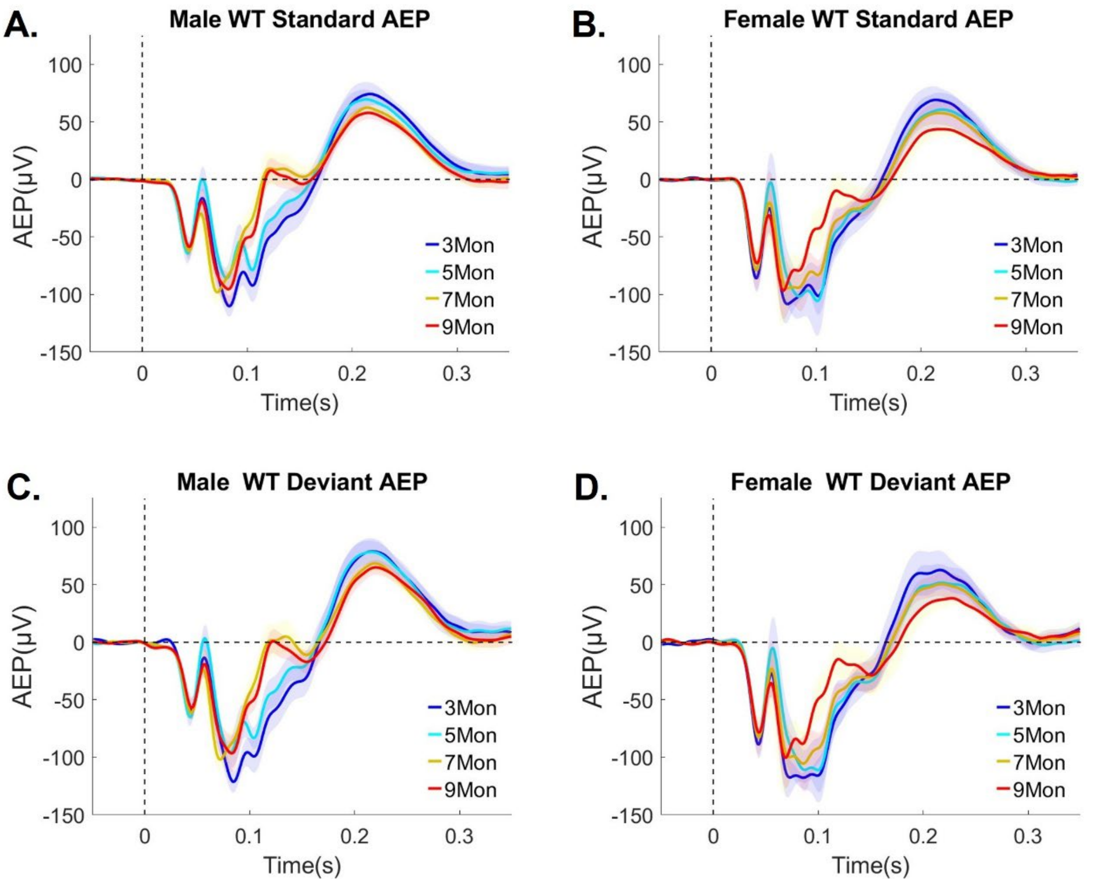

A.
Male WT Standard AEP
B.
Female WT Standard AEP
C.
Male WT Deviant AEP
D.
Female WT Deviant AEP
AEP(μV)
Time(s)
100
50
0
-50
-100
-150
0
0.1
0.2
0.3
3Mon
5Mon
7Mon
9Mon

**Supplementary Fig. 7 WT mice of both sexes showed robust AEP across age at 800 ms ISI.** ***A and B*****,** Trial and subject averaged standard AEP waveforms for male (A) and female (B) WT mice aged from 3 to 9 months, recorded from a centrally located channel (Ch21). The vertical dashed line indicates stimulus onset. ***C and D*****,** Trial and subject averaged deviant AEP waveforms for male (C) and female (D) WT mice aged from 3 to 9 months, recorded from a centrally located channel (Ch21). The vertical dashed line indicates stimulus onset. The shaded areas represent the standard error of the mean (SEM). Blue: 3 month; Cyan: 5 month, Yellow: 7 month, Red: 9 month. Male WT: n = 11 mice for 3-, 7- and 9-month-old groups; n = 14 for the 5-month old group. Female WT: n = 6 mice for all ages. All animals were recorded longitudinally from 3 to 9 months of age, except three additional male mice recorded only at 5 months of age in pilot studies. Male and female WT mice showed robust AEP with age-related change around 100-150 ms time window. The negativity of the AEP around this time window reduced as animal aged. Other peak components also showed slight age-related reduction.

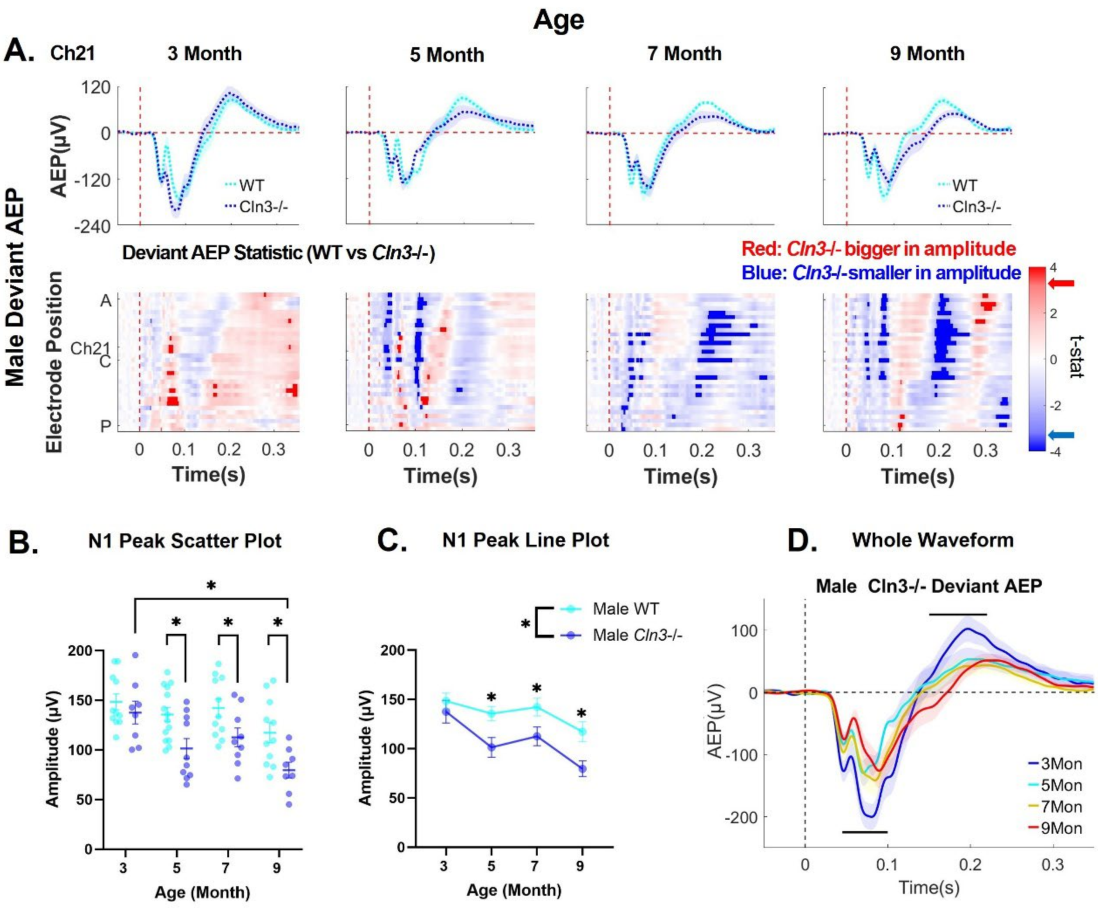

Age
A.
Ch21
3 Month
5 Month
7 Month
9 Month
Male Deviant AEP
AEP(µV)
WT
Cln3-/-
Deviant AEP Statistic (WT vs Cln3-/-)
Red: Cln3-/- bigger in amplitude
Blue: Cln3-/-smaller in amplitude
Electrode Position
t-stat
Time(s)
B.
N1 Peak Scatter Plot
Amplitude (µV)
Age (Month)
C.
N1 Peak Line Plot
Male WT
Male Cln3-/-
D.
Whole Waveform
Male Cln3-/- Deviant AEP
3Mon
5Mon
7Mon
9Mon

**Supplementary Fig. 8 Male *Cln3-/-* mice exhibit progressive reduction of the N1 from 5-9 months.** ***A*,** top row, trial- and subject-averaged deviant AEP waveforms for male WT mice (light blue) and male *Cln3*-/- mice (dark blue) at Ch21. The ISI is 1600 ms. The vertical dashed red line indicates stimulus onset. Bottom row, statistical differences in deviant AEP between male WT and *Cln3*-/- mice were displayed across all electrodes and the entire trial duration (32 electrodes × 400ms). Results from t-tests were corrected for multiple comparisons using the FDR method. Significance was determined when the FDR-adjusted p-value was less than 0.05 (two-tailed). Electrode positions over the mouse skull are indicated from A (anterior) to C (center) and P (posterior). Red indicates electrodes and time bins (in 20ms) where *Cln3*-/- mice showed larger amplitudes (further away from zero compared to WT mice), while blue indicates electrodes and time bins where *Cln3*-/- mice showed smaller amplitudes (closer to zero compared to WT mice). Significant t-stat values are represented by dark red and dark blue, based on the FDR-corrected p-value. Red and blue arrows on the color bar indicate significant t-stat value. Male WT: n = 11 mice for 3-, 7- and 9-month-old groups; n = 14 for the 5-month-old group. Male *Cln3*-/-: n = 8 mice for 3-, 7- and 9-month-old groups; n = 10 for the 5-month- old group. ***B and C*,** scatter and line plots of N1 amplitude (Ch21) for male WT and *Cln3*-/- mice. Individual data points represent each animal and error bar represent mean ± SEM. Repeated measures two-way ANOVA showed a significant genotype effect ($p < 0.0001$, with a progressive reduction in N1 in male *Cln3*-/- mice compared to male WT mice. Age-matched genotype comparison showed that male *Cln3*-/- mice exhibited reductions in N1 at 5, 7 and 9 months of age (3 month: $p = 0.4785$, 5 month: $p = 0.0066$, 7 month: $p = 0.0298$, 9 month: $p = 0.0077$). ***D,*** Trial and subject averaged deviant AEP waveforms for male *Cln3*-/- mice from 3 to 9 months of age (Ch21). The vertical dashed line indicates stimulus onset. Shaded areas represent the standard error of the mean (SEM). Blue: 3 month; Cyan: 5 month, Yellow: 7 month, Red: 9 month. Black horizontal lines indicate the time window during which AEP at 9 months is significantly reduced compared to AEP at 3 months ($p < 0.05$, FDR corrected unpaired t-tests).

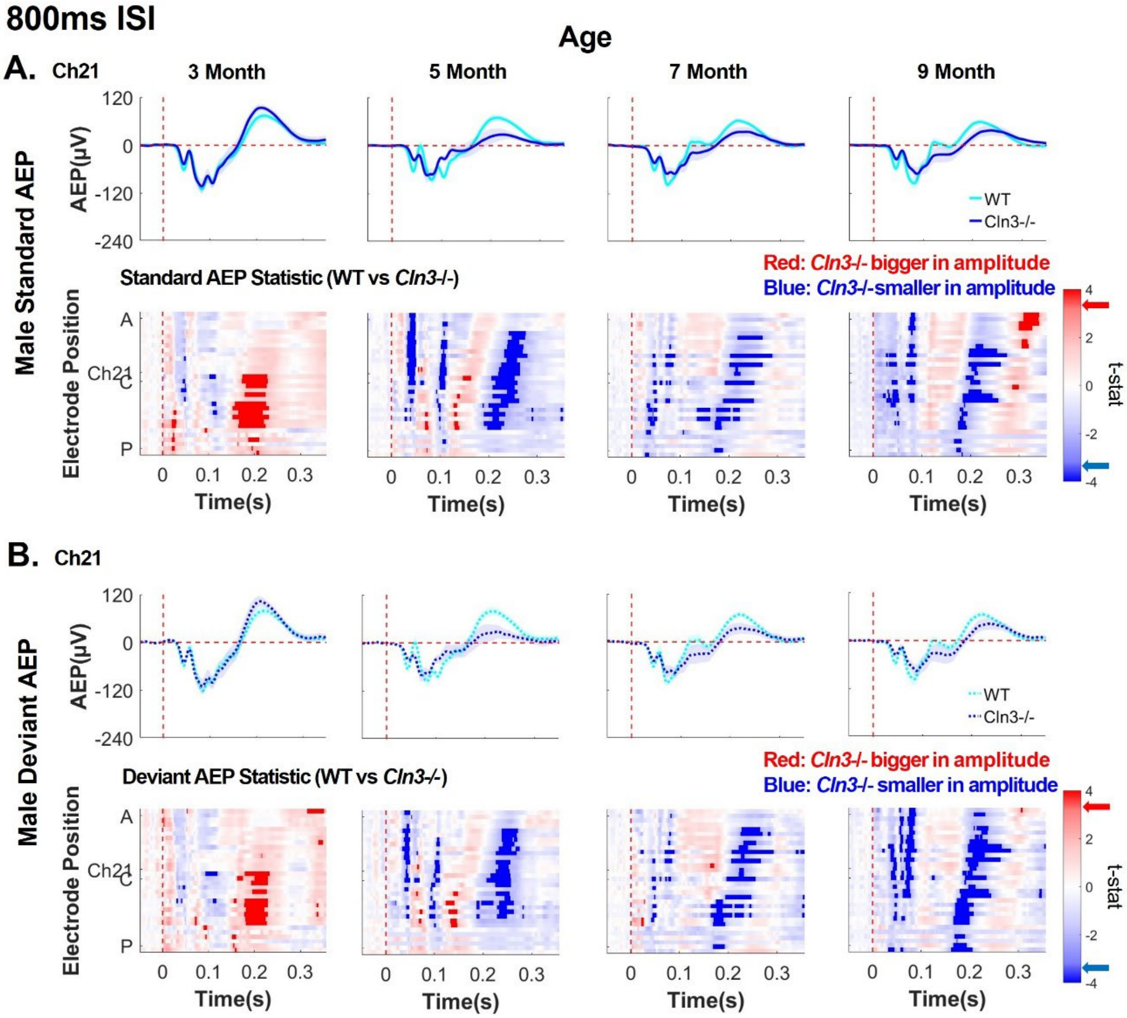

800ms ISI
Age
A.
Ch21
3 Month
5 Month
7 Month
9 Month
Male Standard AEP
AEP(μV)
120
0
-120
-240
WT
Cln3-/-
Standard AEP Statistic (WT vs Cln3-/-)
Red: Cln3-/- bigger in amplitude
Blue: Cln3-/-smaller in amplitude
Electrode Position
A
Ch21
C
P
0
0.1
0.2
0.3
Time(s)
t-stat
4
2
0
-2
-4
B.
Ch21
Male Deviant AEP
Deviant AEP Statistic (WT vs Cln3-/-)
Red: Cln3-/- bigger in amplitude
Blue: Cln3-/- smaller in amplitude

**Supplementary Fig. 9 Male *Cln3-/-* mice exhibited an initial enhancement in AEP followed by a later decline at 800ms ISI.** ***A*,** top row, trial- and subject-averaged standard AEP waveforms for male WT mice (light blue) and male *Cln3*-/- mice (dark blue) at Ch21. The vertical dashed red line indicates stimulus onset. Bottom row, statistical differences in standard AEP between male WT and *Cln3*-/- mice were displayed across all electrodes and the entire trial duration (32 electrodes × 400ms). Results from t-tests were corrected for multiple comparisons using the FDR method. Significance was determined when the FDR-adjusted p-value was less than 0.05 (two-tailed). Electrode positions over the mouse skull are indicated from A (anterior) to C (center) and P (posterior). Red indicates electrodes and time bins (in 20ms) where *Cln3*-/- mice showed larger amplitudes (further away from zero compared to WT mice), while blue indicates electrodes and time bins where *Cln3*-/- mice showed smaller amplitudes (closer to zero compared to WT mice). Significant t-stat values are represented by dark red and dark blue, based on the FDR-corrected p-value. Red and blue arrows on the color bar indicate significant t-stat value. ***B*,** Deviant AEP waveforms at Ch21 (top row) and t-stat values across all electrodes and the entire trial duration (bottom row). Male WT: n = 11 mice for 3-, 7- and 9-month-old groups; n = 14 for the 5-month-old group. Male *Cln3*-/-: n = 8 mice for 3-, 7- and 9-month-old groups; n = 10 for the 5-month- old group.

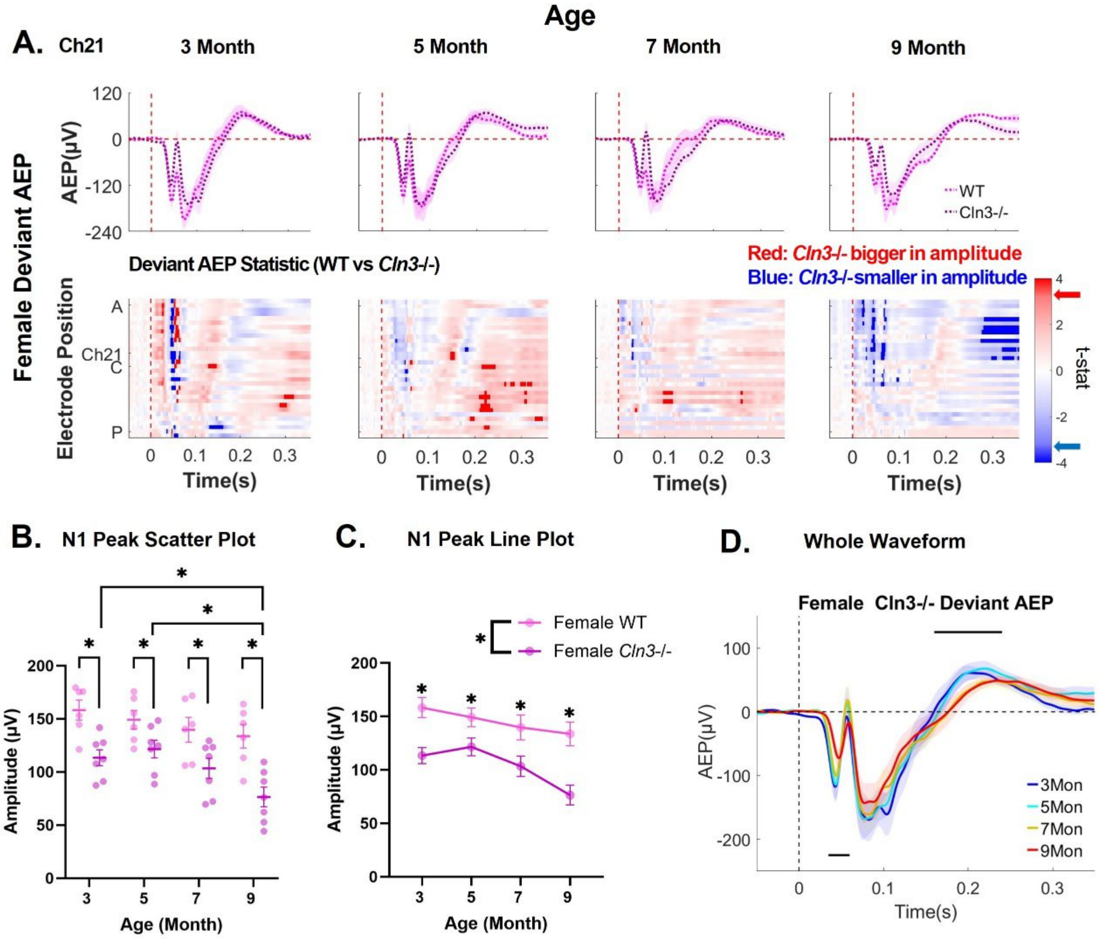

Age
A.
Ch21
3 Month
5 Month
7 Month
9 Month
Female Deviant AEP
AEP(μV)
WT
Cln3-/-
Deviant AEP Statistic (WT vs Cln3-/-)
Red: Cln3-/- bigger in amplitude
Blue: Cln3-/-smaller in amplitude
Electrode Position
t-stat
Time(s)
B.
N1 Peak Scatter Plot
Amplitude (μV)
Age (Month)
C.
N1 Peak Line Plot
Female WT
Female Cln3-/-
Amplitude (μV)
Age (Month)
D.
Whole Waveform
Female Cln3-/- Deviant AEP
AEP(μV)
3Mon
5Mon
7Mon
9Mon
Time(s)

**Supplementary Fig. 10 Female *Cln3-/-* mice exhibit early-onset and progressive reduction of the N1**. ***A,*** top row, trial- and subject-averaged deviant AEP waveforms for female WT mice (light pink) and female *Cln3*-/- mice (dark pink) at Ch21. The ISI is 1600 ms. The vertical dashed red line indicates stimulus onset. Bottom row, statistical differences in deviant AEP between female WT and *Cln3*-/- mice were displayed across all electrodes and the entire trial duration (32 electrodes × 400ms). Results from t-tests were corrected for multiple comparisons using the FDR method. Significance was determined when the FDR-adjusted p-value was less than 0.05 (two-tailed). Electrode positions over the mouse skull are indicated from A (anterior) to C (center) and P (posterior). Red indicates electrodes and time bins (in 20ms) where *Cln3*-/- mice showed larger amplitudes (further away from zero compared to WT mice), while blue indicates electrodes and time bins where *Cln3*-/- mice showed smaller amplitudes (closer to zero compared to WT mice). Significant t-stat values are represented by dark red and dark blue, based on the FDR-corrected p-value. Red and blue arrows on the color bar indicate significant t-stat value. Female WT: $n = 6$. Female *Cln3*-/-: $n = 7$. ***B and C,*** deviant N1 amplitude (Ch21) scatter and line plots for female WT and *Cln3*-/- mice. Individual data points for each animal and mean ± SEM. Repeated measures two-way ANOVA showed a significant genotype effect ($p < 0.0001$, with a progressive reduction in N1 in female *Cln3*-/- mice compared to female WT mice. Age-matched genotype comparison showed that female *Cln3*-/- mice exhibited reductions in N1 across age (3 month: $p = 0.0021$, 5 month: $p = 0.0479$, 7 month: $p = 0.0117$, 9 month: p = 0.0001). ***D,*** Trial and subject averaged deviant AEP waveforms for female *Cln3*-/- mice from 3 to 9 months of age (Ch21). The vertical dashed line indicates stimulus onset. Shaded areas represent the standard error of the mean (SEM). Blue: 3 month; Cyan: 5 month, Yellow: 7 month, Red: 9 month. Black horizontal line indicates the time window during which AEP at 9 months is significantly reduced compared to AEP at 3 months ($p < 0.05$, FDR corrected unpaired t-tests).

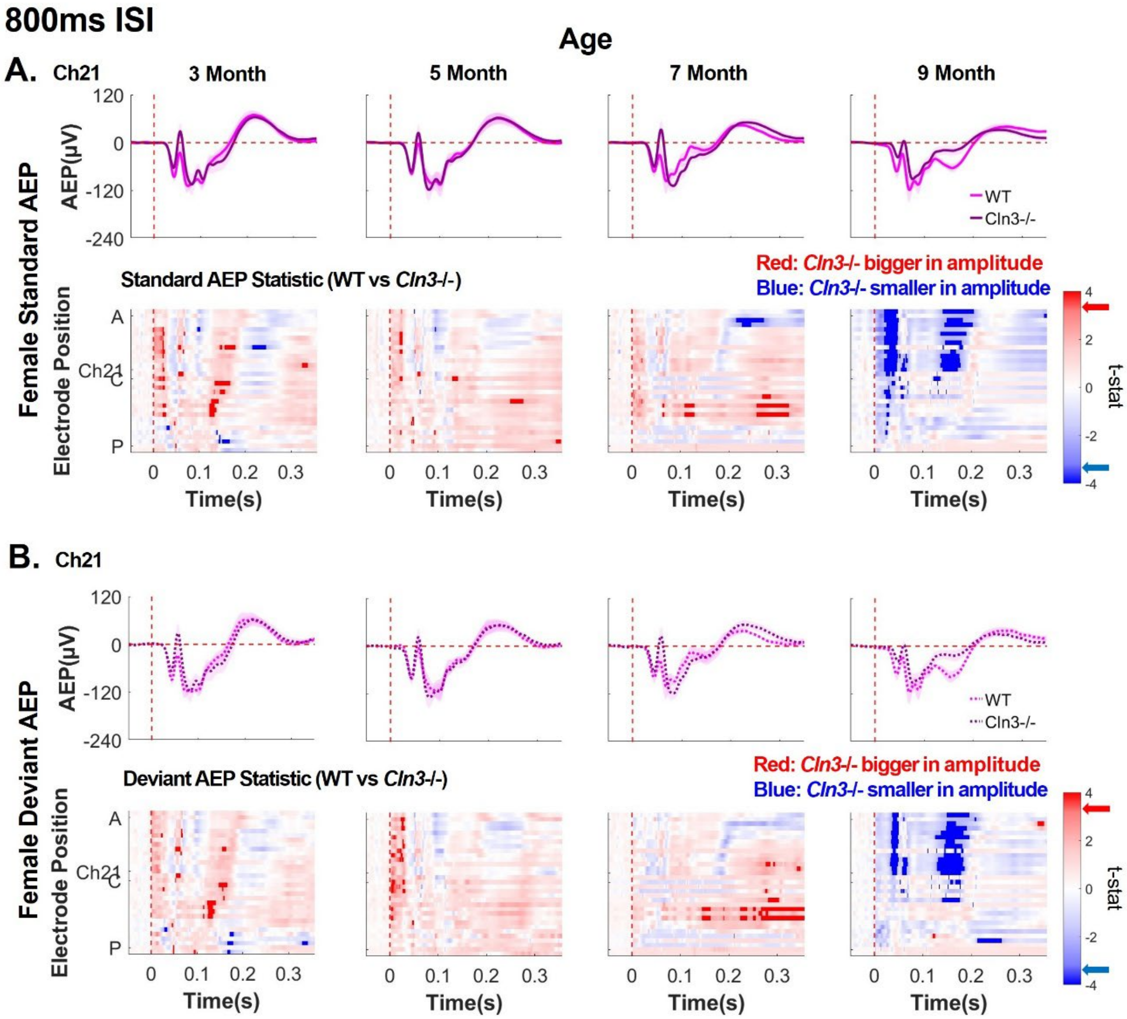
800ms ISI
Age
A.
Ch21
3 Month
5 Month
7 Month
9 Month
Female Standard AEP
AEP(µV)
WT
Cln3-/-
Standard AEP Statistic (WT vs Cln3-/-)
Red: Cln3-/- bigger in amplitude
Blue: Cln3-/- smaller in amplitude
Electrode Position
Time(s)
t-stat
B.
Ch21
Female Deviant AEP
Deviant AEP Statistic (WT vs Cln3-/-)
Red: Cln3-/- bigger in amplitude
Blue: Cln3-/- smaller in amplitude

**Supplementary Fig. 11 Female *Cln3*-/- showed pronounced reductions in AEPs at 9 months of age at 800ms ISI.** ***A*,** top row, trial- and subject-averaged standard AEP waveforms for female WT mice (light pink) and male *Cln3*-/- mice (dark pink) at Ch21. The vertical dashed red line indicates stimulus onset. Bottom row, statistical differences in standard AEP between female WT and *Cln3*-/- mice were displayed across all electrodes and the entire trial duration (32 electrodes × 400ms). Results from t-tests were corrected for multiple comparisons using the FDR method. Significance was determined when the FDR-adjusted p-value was less than 0.05 (two-tailed). Electrode positions over the mouse skull are indicated from A (anterior) to C (center) and P (posterior). Red indicates electrodes and time bins (in 20ms) where *Cln3*-/- mice showed larger amplitudes (further away from zero compared to WT mice), while blue indicates electrodes and time bins where *Cln3*-/- mice showed smaller amplitudes (closer to zero compared to WT mice). Significant t-stat values are represented by dark red and dark blue, based on the FDR-corrected p-value. Red and blue arrows on the color bar indicate significant t-stat value. ***B*,** Deviant AEP waveforms at Ch21 (top row) and t-stat values across all electrodes and the entire trial duration (bottom row). Female WT: n = 6. Female *Cln3*-/-: n = 7. Female *Cln3*-/- mice showed pronounced reductions in deviant AEPs at 9 months of age with persistent decrease in N1.
